# Endogenous auditory and motor brain rhythms predict individual speech tracking

**DOI:** 10.1101/2025.03.24.644939

**Authors:** Christina Lubinus, Anne Keitel, Jonas Obleser, David Poeppel, Johanna M. Rimmele

**Affiliations:** Department of Cognitive Neuropsychology, Max-Planck-Institute for Empirical Aesthetics, 60322 Frankfurt am Main, Germany; Psychology, University of Dundee, Dundee DD1 4HN, UK; Department of Psychology, University of Lübeck, Lübeck, Germany; Center for Brain, Behavior, and Metabolism, University of Lübeck, Lübeck, Germany; Department of Psychology, New York University, New York, NY, USA

**Author notes:** Corresponding author: Johanna M. Rimmele.

## Abstract

Slow, endogenous brain rhythms in auditory cortex are hypothesized to track acoustic amplitude modulations during speech comprehension. Temporal predictions from the motor system are thought to enhance this tracking. However, direct evidence for the involvement of endogenous auditory and motor brain rhythms is lacking. Combining magnetoencephalographic recordings with behavioral data, we here show that endogenous peak frequencies of individuals’ resting-state theta rhythm in superior temporal gyrus predict speech tracking during comprehension. Importantly, endogenous rates of speech motor areas predicted auditory-cortical speech tracking only in individuals with high auditory–motor synchronization profiles. Higher rates in the supplementary motor area, and lower rates in inferior frontal gyrus, predicted stronger tracking. These findings align with participants’ behavioral data and provide compelling support for oscillatory accounts of auditory–motor interactions during speech perception. Interestingly, working memory capacity predicted speech comprehension performance particularly in individuals with low auditory–motor synchronization profiles. The findings highlight two partially independent speech processing routes across individuals: an auditory–motor route, related to enhanced comprehension performance, and an auditory working-memory route.

## Introduction

Verbal communication exemplifies action-perception interactions in humans essential for everyday behavior. During speech production, the motor system engages in operations requiring precise timing (1,2). The speech motor system may also be recruited during speech perception (1,3,4), with the supplementary motor area (SMA) and inferior frontal gyrus (IFG) generating temporal predictions about upcoming sensory events (5–10). A prominent neural oscillatory account of speech perception proposes that slow endogenous brain rhythms in auditory cortex allow for speech segmentation by aligning their neural excitability phase to the speech acoustics (speech tracking; ,11–16). Additionally, neural oscillations from motor cortices may be involved in speech perception. Slow and fast neural oscillations supposedly aid temporal predictions from motor cortices through coupling with auditory areas (5,6,17,18). The mechanisms of auditory-motor interactions during speech (and auditory) perception, however, are not fully understood and the involvement of neural oscillations is controversially discussed (19–21). A crucial characteristic of neural oscillations is that they reflect endogenous brain rhythms observed in the absence of external stimulation. Here, we put a neural oscillatory framework of auditory-motor interactions to a rigorous test by investigating whether individuals’ peak frequencies of endogenous rhythms of the auditory and motor systems (observed during resting-state) and their coupling strength predict auditory cortical tracking during speech comprehension.

Oscillatory speech perception models propose that endogenous theta rhythms in auditory cortex synchronize to temporal fluctuations in the speech signal (i.e. the amplitude envelope) to segment it into syllable-sized chunks (12–15,22,23). This brain-to-speech alignment is most pronounced in the theta range (∼5Hz), declining at higher syllabic rates (24), as speech comprehension also decreases (for non-speech see: ,25–29). Given the observation of endogenous theta brain rhythms in auditory cortex (30–32) and the optimal speech processing in this range, the preferred frequencies of neuronal populations in auditory cortex in the theta range have been proposed to constrain the temporal granularity of perception (14,22,32–36). The hypothesized connection between endogenous theta rhythms and speech processing, a fundamental aspect of oscillatory theories, however, has been rarely investigated directly, i.e. by relating endogenous and functional processing within individuals (37). Such research may be hindered by the difficulty of quantifying individual endogenous brain rhythms in auditory cortex in the theta range (34). Spectral fingerprinting of resting-state brain activity (34) uses spectral clustering across time and a normalization procedure, which makes it a promising approach for accessing individual differences in the rates of endogenous theta rhythms.

According to recent work, not only auditory processing but also auditory-motor coupling has an optimal range (∼4.5Hz; Assaneo and Poeppel, 2018; He et al., 2023), hinting at the involvement of motor cortex oscillations in the auditory-motor interaction. Computational modelling supports an oscillatory account of auditory-motor coupling during speech perception, assuming oscillators with slightly higher preferred frequencies in the auditory than the motor system (17,39). Behavioral and computational studies have shown a relation between preferred (spontaneous) motor production rates and the ability to synchronize to sound at different rates in music and sound sequences (for a neural approach see: ,40–43). In the context of dynamical systems theory, the *Arnold tongue* phenomenon describes how an oscillator’s ability to track a stimulus depends on its preferred frequency and the frequency and intensity of an external stimulation (Fröhlich & McCormick, 2010). However, how this phenomenon transfers to the preferred frequencies of two oscillatory systems (auditory and motor) and their coupling strength to shape our ability to track the speech acoustics is unknown. In a recent behavioral study (29), we approached this question, demonstrating superior speech comprehension in individuals with higher auditory-motor synchronization and higher preferred motor rates. The spontaneous speech synchronization (SSS) test was used to behaviorally estimate auditory-motor cortex coupling strength (44). The preferred endogenous motor rates were quantified by the spontaneous rhythmic speech production rates of an individual, like the spontaneous tapping measure typically used in the field of dynamic attending (41,45–49).

Here, we use a behavioral and MEG approach in a relatively large sample, combining spectral fingerprinting of individuals’ MEG resting-state brain activity with MEG data recorded during a speech comprehension task (Fig. 3 A), and with additional behavioral tasks (Fig. 3 A,C,E; and the digit-span to access working memory capacity). First, we aim to test the neural oscillatory approach of speech segmentation by quantifying predictive effects of individual endogenous theta rates in auditory cortex on speech tracking and comprehension. Second, to characterize the conjectured neural oscillatory approach of auditory-motor interactions, we expect predictive effects of individual theta rates in motor cortices and the auditory-motor coupling strength. Our behavioral findings replicate and further specify our previous study (29), by showing that higher spontaneous preferred auditory rates and auditory-motor synchronization predict speech comprehension. In contrast, higher spontaneous speech motor production rates only predicted comprehension in parts of the population (high audio-motor synchronizers), with low synchronizers instead showing predictive effects of their working memory capacity. Importantly, the neural results parallel the behavioral findings. Only in individuals with high auditory–motor synchronization profiles, the endogenous rates of speech motor cortices in SMA and IFG predicted speech tracking in auditory cortices, STG and HG, respectively. In contrast, endogenous auditory rates of STG (but not HG) were predictive of speech tracking across the population. We provide compelling evidence for a neural oscillatory account that highlights two distinct speech processing routes across individuals.

## Results

### Speech comprehension predicted by behavioral auditory and motor parameters

In a behavioral session, participants (N = 57) completed behavioral tasks assessing their spontaneous speech motor production rate (henceforth *motor production rate*) (Fig 1C), preferred auditory rate (Fig 1E), and auditory-motor synchronization (Fig 1A). Participants exhibited an average motor production rate of M = 4.24 syllables/s (SD = 0.51, Fig 1D). Their preferred auditory rate was about one syllable/s faster than the motor rate with the average at M = 5.61 syllables/s (SD = 0.78, Fig 1F). Consistent with previous work (29,39,44), auditory-motor synchronization, measured using the SSS-test, was consistent with a bimodal distribution (Fig 1B). In this sample, N = 27 participants were classified as high synchronizers and N = 20 as low synchronizers (N = 10 participants were not clearly classified at probabilities of being high synchronizers around ∼50% and thus were excluded from the behavioral analyses).

**Figure 1.**
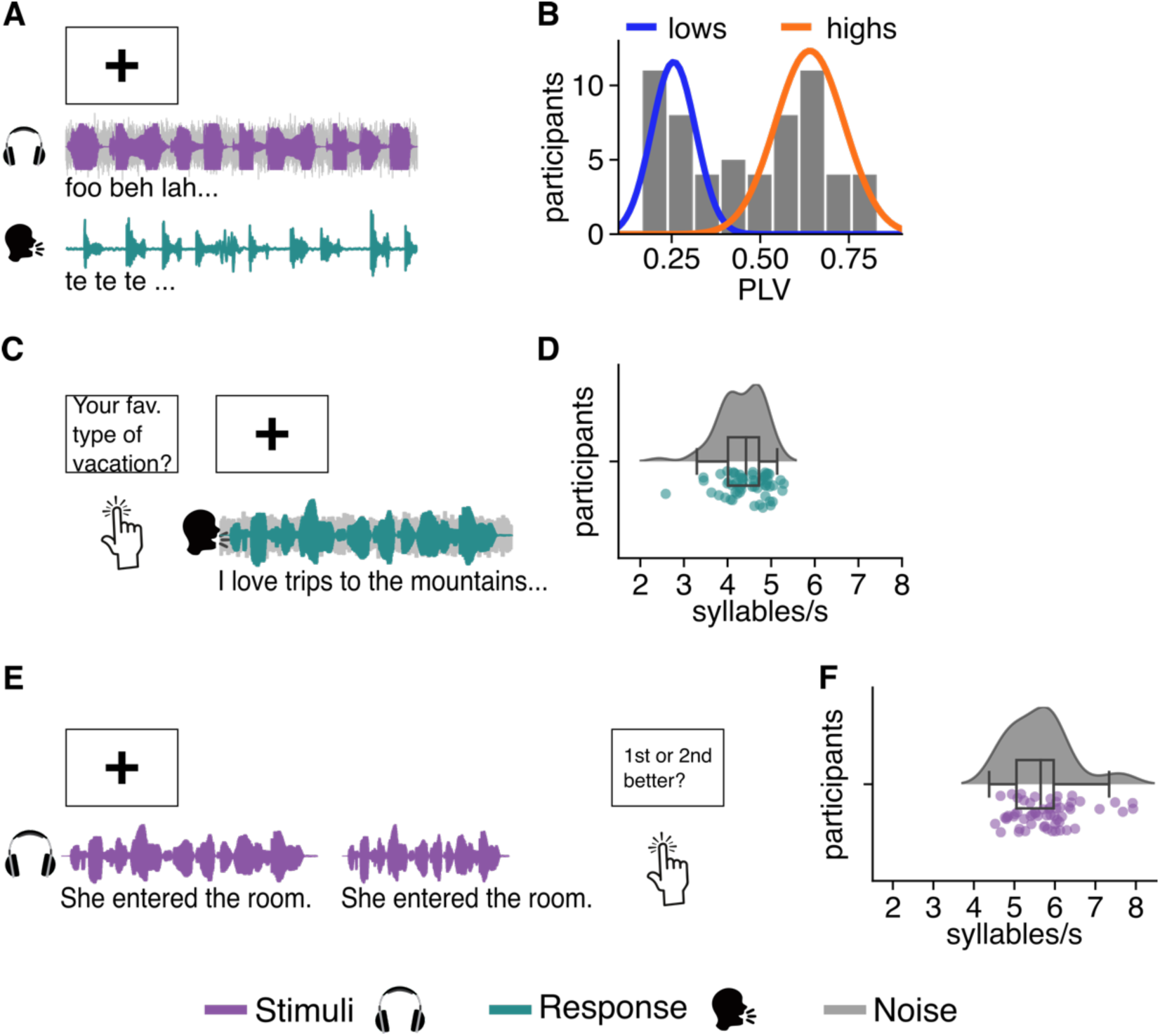
Experimental paradigms and behavioral data. **A.** We measured auditory-motor synchronization using the (explicit) SSS-test (50), wherein participants whisper the syllable /te/ while listening to a syllable train. They are instructed to synchronize their motor output to the auditory input. **B.** To quantify the motor rate, we conducted a speech motor production task. Upon being prompted with a question/statement, participants spoke freely for 30s. **C.** In the preferred auditory rate preference task, participants indicated which speech rates they liked better. To this end, two versions of a sentence, differing in their syllabic rates, were presented on each trial and participants chose the sentence (i.e. syllabic rate) they preferred. **D.** Histogram illustrates audio-motor synchronization (measured as PLV). Colored lines represent fitted normal distribution, obtained by a Gaussian mixture model. **E, F.** Density and dot plots visualize the motor production rate (**E**) and preferred auditory rate (**F**) of participants.

Speech comprehension was measured using an intelligibility task during the MEG recording session (Fig 3A,C). Trial-based comprehension accuracy (% words correct) was regressed against several variables (see Fig 2A, B and Supplementary Table 3). FDR corrected p-values are reported, if not otherwise indicated. As expected, we observed a main effect of syllabic rate, with decreased comprehension for higher syllabic rates (linear: β = −70.81, SE = 1.78, p_FDR_ < .001; quadratic: β = −24.36, SE = 0.98, p_FDR_ < .001). Replicating previous findings (29), stronger auditory-motor synchronization (β = 0.03, SE = 0.02, p_FDR_ = .032), and faster motor production rates (β = 0.03, SE = 0.02, p = .044, not FDR-corrected) predicted better speech comprehension. In contrast to our previous findings, the motor production rate effect did not survive the control for multiple comparisons (FDR-corrected: p_FDR_ = .056). In addition, faster preferred auditory rates predicted better speech comprehension (β = 0.09, SE = 0.02, p_FDR_ < .001). These findings were refined by two-way interaction effects of PLV ✕ motor production rate (β = 0.07, SE = 0.02, p_FDR_ < .001) and motor production rate ✕ preferred auditory rate (β = −0.06, SE = 0.02, p_FDR_ = .002), and a three-way interaction of motor production rate ✕ preferred auditory rate ✕ PLV (β = −0.06, SE = 0.02, p_FDR_ < .001). The positive effect of the motor production rate was particularly observed in high synchronizers (PLV ✕ motor production rate), whereas the positive effect of the preferred auditory rate was stronger in individuals with low motor production rates. The 3-way interaction suggests that the interplay of auditory and motor rates is particularly evident in individuals with stronger auditory-motor synchronization. Several control variables facilitated speech comprehension: better working memory performance (β = 0.07, SE = 0.01, p_FDR_ < .001), shorter sentences (β = −0.11, SE = 0.02, p_FDR_ < .001) and occurrence late in the experiment (β = 0.05, SE = 0.00, p_FDR_ < .001).

**Figure 2.**
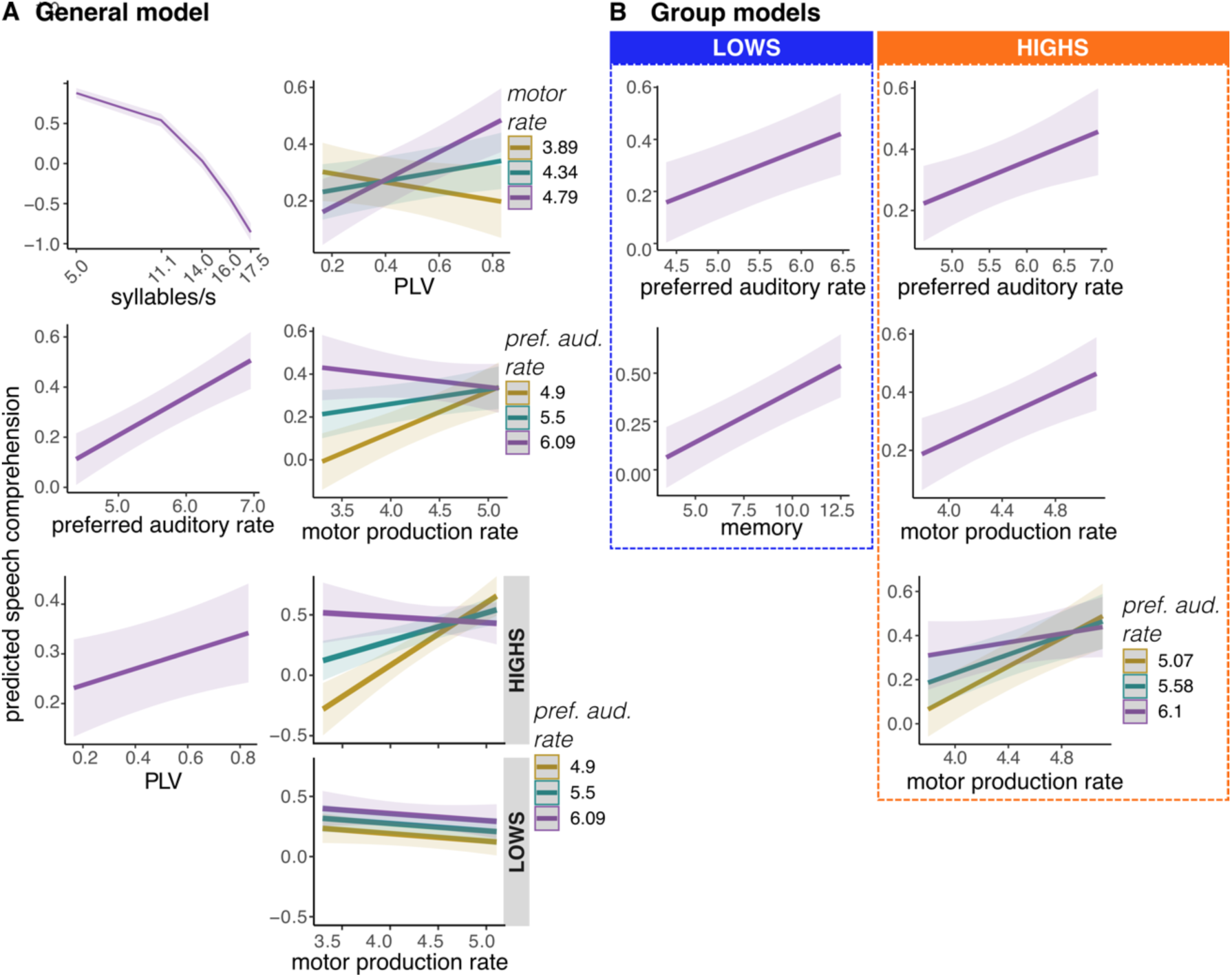
Main and interaction effects predicting speech comprehension in all participants (A) and in high versus low synchronizers (B). **A.** The general GLMM revealed a negative main effect of syllabic rate (left column, row 1) and positive main effects of the preferred auditory rate (left column, row 2) and PLV (left column, row 3). We further observed interaction effects (right column) between preferred motor rate and PLV (right column, row 1), preferred auditory rate and motor production rate (right column, row 2), and a three-way interaction of preferred auditory rate, motor production rate and PLV (right column, row 3-4) **B.** Results from GLMMs computed separately for low (blue, left column) and high (orange, right column) synchronizers. In low synchronizers, speech comprehension was predicted by the preferred auditory rate (left column, row 1) and working memory (left column, row 2). In contrast, for high synchronizers preferred auditory rate (right column, row 1), motor production rate (right column, row 2), and their interaction (right column, row 3) predicted speech comprehension.

Because it has been hypothesized that high and low synchronizers behave fundamentally differently (39) and to reduce the complexity of the model, we computed separate models for the two groups (Fig 2B and Supplementary Table 4). Both groups showed a main effect of syllabic rate (LOW: (linear: β = −45.67, SE = 2.02, p_FDR_ < .001; quadratic: β = −15.82, SE = 0.92, p_FDR_ < .001); HIGH: (linear: β = −53.61, SE = 1.69, p_FDR_ < .001; quadratic: β = −19.73, SE = 0.98, p_FDR_ < .001)) and a main effect of preferred auditory rate (LOW: (β = 0.08, SE = 0.02, p_FDR_ < .001), HIGH: (β = 0.05, SE = 0.02, p_FDR_ = .010). For low synchronizers, we further observed an effect of working memory (β = 0.13, SE = 0.02, p_FDR_ < .001). In high synchronizers, speech perception was additionally predicted by the motor production rate (β = 0.08, SE = 0.02, p_FDR_ < .001), and a motor production rate ✕ preferred auditory rate interaction effect (β = −0.04, SE = 0.02, p_FDR_ = .013). While the statistical power was lower in the model of low synchronizers due to fewer participants (N=20 vs. N=27), these results suggest—jointly with the three-way interaction of the main model—that the motor production rate effects and the interaction of the auditory and motor rates are mainly present in individuals with high auditory-motor synchronization.

### Auditory cortex tracking and auditory-motor coupling across syllabic rates

Next, we analyzed whether speech tracking could be predicted by auditory-motor coupling and the endogenous theta frequencies of auditory (HG, STG) and speech motor (IFG, SMA) brain areas. To quantify speech tracking (see Fig 3B for analysis pipeline), we computed Gaussian-Copula Mutual Information (GCMI; 51) between neuronal activity in auditory brain areas (HG and STG) and the speech signal’s amplitude envelope. To test for significant tracking, GCMI was normalized using surrogate data, yielding z-transformed GCMI values (see Supplementary Fig 1). Speech tracking peaked at the syllabic rate of sentences relative to the other tested frequencies (Fig. 3E).

**Figure 3.**
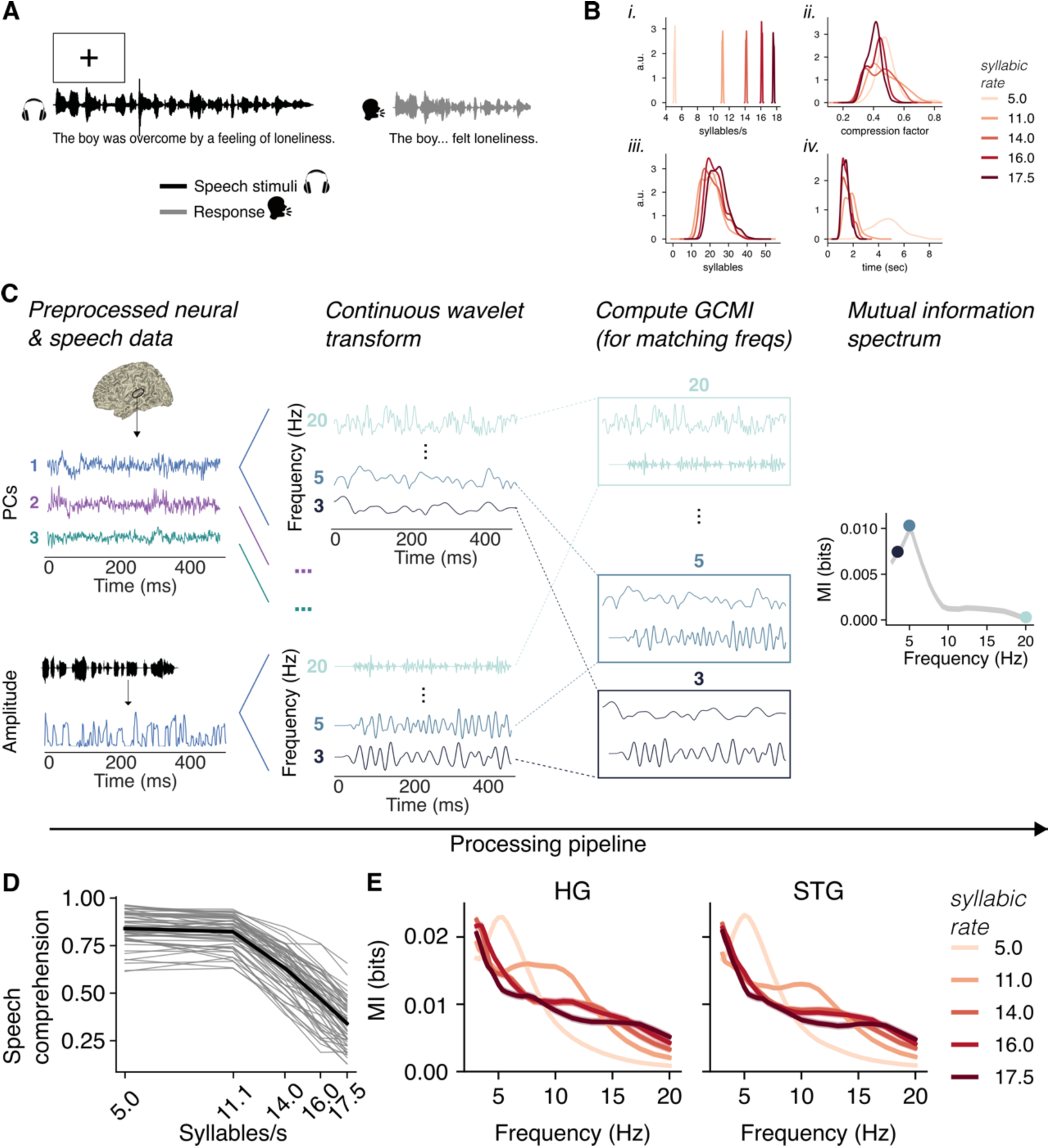
Speech tracking in HG and STG at all syllabic rates. **A.** During the MEG recording session, participants performed the intelligibility task. **B.** Stimulus parameters for sentences presented in the comprehension task. *i.* Sentences were presented at five syllabic rates: 5.0, 11.0, 14.0, 16.0, 17.5 syllables/s. *ii.* Compression differences were minimized between rate conditions. *iii.* Information density across rate conditions was balanced by selecting sentences with overlapping distributions of sentence length (i.e. number of syllables). *iv.* However, this came at the cost of not fully equalizing duration. While the four faster conditions were similar in duration, sentences in the slowest condition (5 syllables/s) were notably longer. **C.** Illustration of Gaussian-Copula Mutual Information (GCMI) processing pipeline (reads from left to right). **D.** Line graph displays the behavioral results of intelligibility task: speech was comprehended well at 5 and 11 syllables/s; comprehension deteriorated for the higher syllabic rates. Grey lines illustrate single participant comprehension, thick black line represents the mean over participants. **E.** Non-normalized GCMI spectra in HG and STG. For both panels, lines are color-coded according to the syllabic rate of sentence stimuli (see legend) and shaded error bars represent standard error of the mean across participants. HG = Heschl’s gyrus, STG = superior temporal gyrus; Next, we quantified auditory-motor coupling by computing GCMI between speech motor areas (IFG tri. and SMA) and auditory areas (HG and STG; see Supplementary Fig 2A). Using the optimal delay per condition, we extracted GCMI spectra for all conditions and ROI pairs. GCMI was again normalized using surrogate data, yielding z-transformed GCMI values. However, the normalized GCMI spectrum showed an offset in the 5 syllables/s condition relative to the faster conditions (see Supplementary Fig 2B). Consequently, we treat the normalized data cautiously and conduct subsequent analyses on the non-normalized GCMI values for both speech tracking and auditory-motor coupling.

### Individual rate of endogenous theta brain rhythms in auditory and motor cortex

We hypothesized that the individual rate of the endogenous theta rhythms in auditory and speech motor areas (‘eigenfrequency’, as assessed during resting-state) predicts speech tracking. To test this hypothesis, we identified spectral fingerprints in HG, STG, IFG, and SMA using a clustering approach (see Supplementary Fig 3). From single-subject clusters, we extracted each participant’s theta frequency for these brain regions (Fig 4A,C). While not all subjects exhibited a theta cluster in every ROI, the majority of subjects did (Fig 5B; HG-L: N = 55; HG-R: N = 56, STG-L: N = 56, STG-R: N = 55, IFG-L: N = 56, IFG-R: N = 56, SMA-L: N = 57, SMA-R: N = 56).

**Figure 4.**
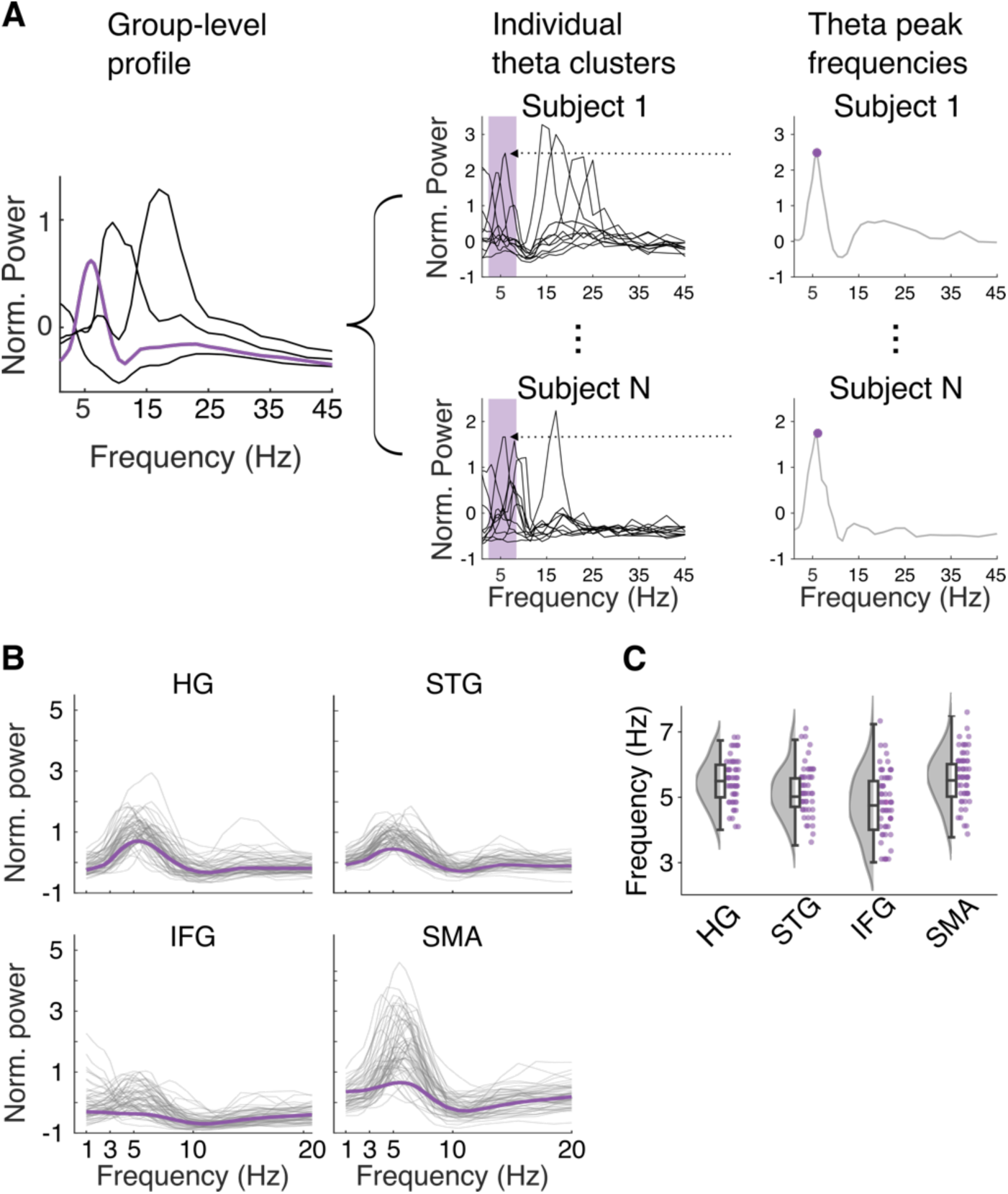
Spectral fingerprints of HG, STG, IFG, and SMA. **A.** Illustration of single-subject theta cluster extraction. Group-level spectral profiles contain multiple spectral clusters, each represented by a line, express region-specific spectral power relative to the power across the whole brain. For further analysis only the cluster peaking in the theta range (i.e. magenta line) was considered. We reconstructed which single-subject clusters (multiple clusters possible) contributed to the group-level theta cluster and selected the individual theta cluster with the highest amplitude. Finally, the frequency at the cluster peak was extracted as individual theta frequency. **B.** Line graphs illustrate individual theta clusters (grey lines) corresponding to the group-level clusters (thick magenta line), averaged across hemispheres. Note that for IFG the group cluster is also colored in magenta (not grey as done above in panel E) to better distinguish group and individual clusters. **C.** Distribution- and boxplots visualize the peak frequencies extracted from the individual theta clusters. HG = Heschl’s gyrus; STG = superior temporal gyrus; IFG = interior frontal gyrus; SMA = supplementary motor area.

**Figure 5.**
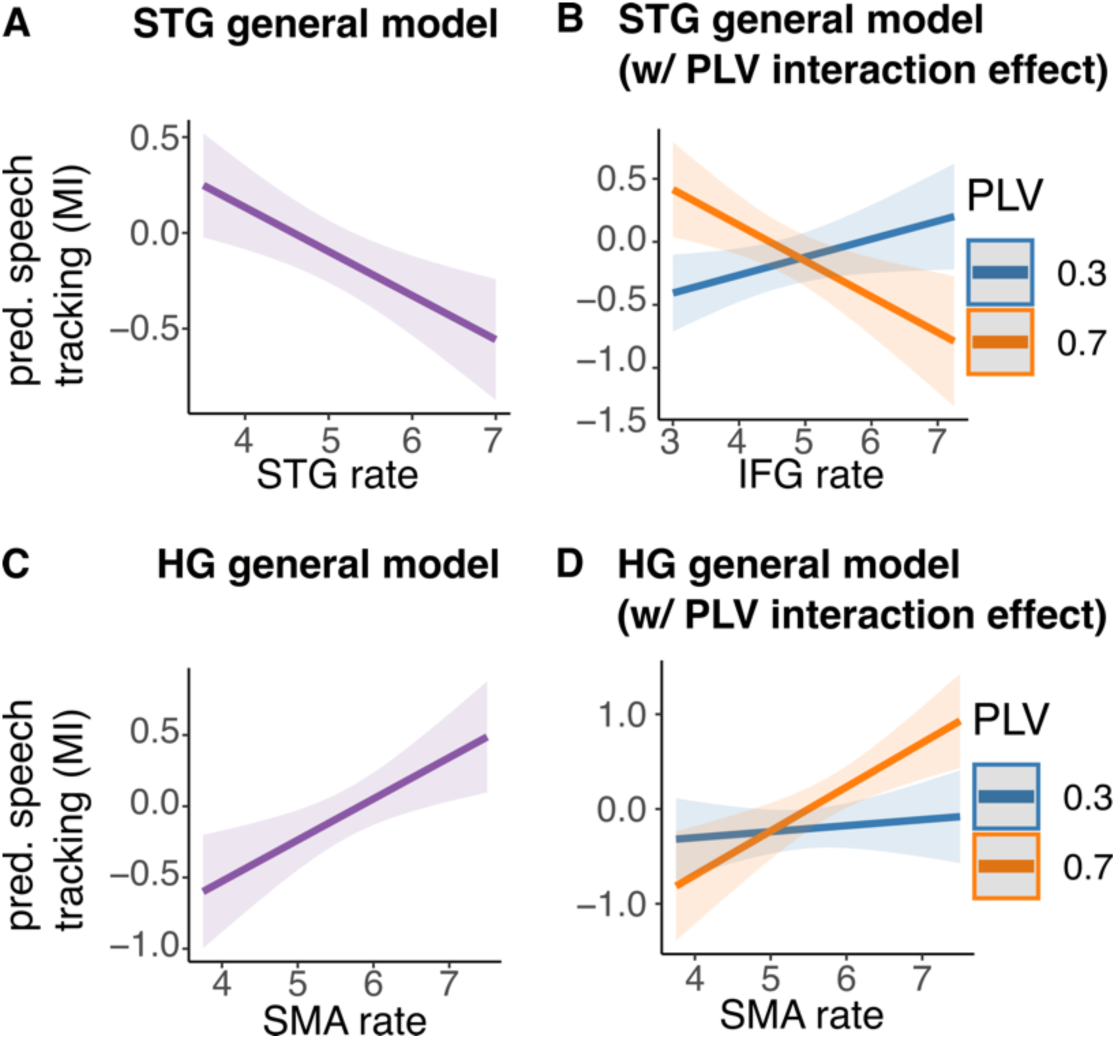
Speech tracking in STG and HG is predicted by distinct auditory and motor variables. **A.** In the STG general model, we observed a negative main effect of the endogenous rate of STG on speech tracking (in STG). **B.** Including two-way interactions between the behavioral measure of auditory-motor synchronization and motor brain rhythms revealed a significant interaction effect of PLV ✕ IFG, but not PLV ✕ SMA, on speech tracking in STG. **C.** The HG general model revealed a main effect of SMA frequency on speech tracking in HG. **D.** In the HG model with PLV interaction terms we observed an interaction effect of PLV ✕ SMA, but not of PLV ✕ IFG, on speech tracking. In all panels, error shades indicate 95% confidence intervals. HG = Heschl’s gyrus, STG = superior temporal gyrus; IFG = interior frontal gyrus; SMA = supplementary motor area.

### Auditory-motor parameters affect speech tracking in HG and STG differentially

We combined all neural variables to assess whether speech tracking was affected by the endogenous theta frequency of auditory (HG, STG) and speech motor (IFG, SMA) brain areas, as well as auditory-motor coupling. To this end, we computed two GLMMs, a “HG general model” and an “STG general model”. FDR corrected p-values are reported, if not otherwise indicated.

The STG general model (N = 50; Fig 5A, Supplementary Table 8) revealed a negative main effect of syllabic rate (linear: β = −8.75, SE = 0.83, p_FDR_ < .001), reflecting a decrease in speech tracking in STG with increasing syllabic rate. Importantly, the model also showed a main effect of the endogenous theta frequency of STG (β = −0.18, SE = 0.06, p_FDR_ = .011), suggesting increased speech tracking for individuals with lower endogenous auditory theta frequencies.

The HG general model (N = 54, Fig 5C, Supplementary Table 5) also revealed a negative main effect of syllabic rate (linear: β = −9.27, SE = 0.80, p_FDR_ < .001). For the endogenous theta frequency of SMA, we observed a positive main effect (β = 0.22, SE = 0.07, p_FDR_ = .023), such that speech tracking was higher in individuals with higher endogenous theta rates in SMA. We further observed a three-way interaction of theta frequency in HG ✕ theta frequency in IFG ✕ IFG-coupling (β = 0.19, SE = 0.08, p = .021), suggesting that IFG coupling and endogenous theta frequency in IFG interact more strongly in individuals with lower endogenous frequencies in HG. The interaction effect, however, did not remain significant after FDR correction (p_FDR_ = 0.094).

### Motor parameters predict speech tracking in high synchronizers

Based on theoretical assumptions and the behavioral findings, we included two-way interaction effects of PLV and both speech motor rates (IFG, SMA) in the HG and STG general models (see Figure 5 B,D; Supplementary Tables 6 and 9). In the STG model (N = 50), we observed a significant interaction effect of PLV and IFG (β = −1.01, SE = 0.28, p_FDR_ = .003, p < .001), but not PLV and SMA (β = 0.32, SE = 0.43, SE = 0.27, p = .281, p_FDR_ = .118). In contrast, the HG model (N = 54) revealed an interaction effect of PLV and SMA (β = 0.77, SE = 0.29, p =.009, p_FDR_ = .089), but not PLV and IFG ( β = −0.62, SE = 0.33, p = .059, p_FDR_ = .224). As both interaction effects suggest that speech tracking is influenced by the motor rate more strongly in individuals with higher auditory-motor synchronization, we also computed separate GLMMs for high and low synchronizers (see Supplementary Tables 7 and 10).

In low synchronizers, the HG model (N = 20) revealed an interaction effect of the endogenous theta frequencies of IFG and HG (β = −0.69, SE = 0.18, p_FDR_ = .002, Fig 6B) with speech tracking being highest when IFG and HG theta frequencies were mismatching (i.e. high IFG and low HG frequency, Fig 6A). In the low synchronizers STG model (N = 19), we observed a negative main effect of endogenous theta frequency in STG (β = −0.49, SE = 0.14, p_FDR_ = .008; Fig 6A).

**Figure 6.**
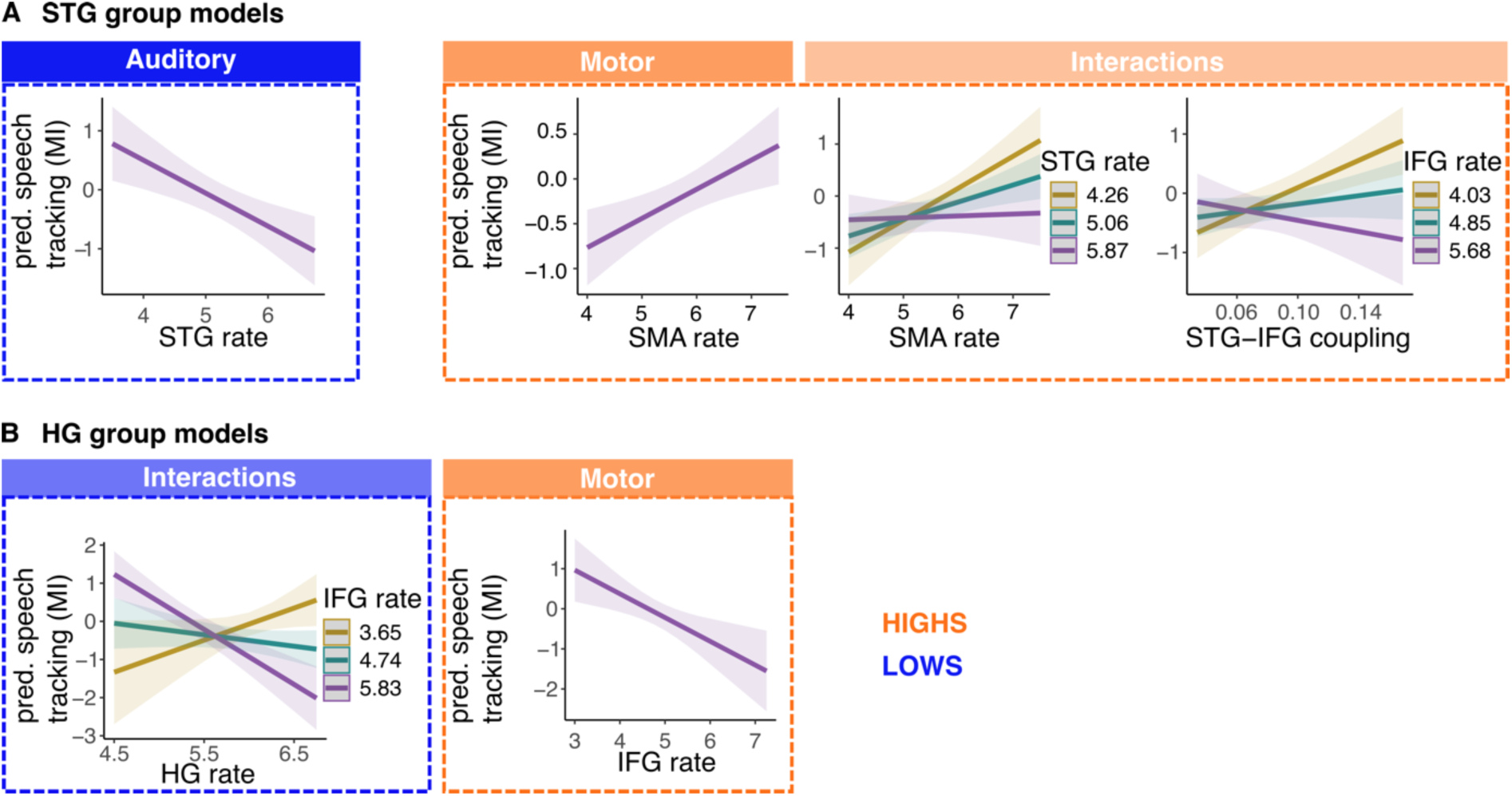
Speech tracking in HG and STG is predicted by different variables in low (blue) and high (orange) synchronizers. **A.** For the STG model, we observed a negative main effect of STG frequency in the low synchronizers. In the high synchronizers, several effects involving speech motor areas were observed: a positive main effect of the endogenous SMA frequencies as well as interaction effects of STG ✕ SMA rates, and STG-IFG coupling ✕ IFG rate. **B.** In low synchronizers, the HG model revealed an interaction effect of the endogenous frequencies of HG and IFG. In high synchronizers, we observed a negative main effect of the endogenous IFG rate. In all panels, error shades indicate 95% confidence intervals. HG = Heschl’s gyrus, STG = superior temporal gyrus; IFG = interior frontal gyrus; SMA = supplementary motor area.

In high synchronizers, the HG model (N = 27, Fig 6B) showed a negative main effect for endogenous theta frequency in IFG (β = −0.44, SE = 0.14, p_FDR_ = .025). Other effects were observed, however, did not survive correction for multiple comparisons (Supplementary Table 7). The high synchronizers’ STG model (N = 27, Fig 6A) failed to converge. Therefore, we recomputed the model using a simplified random effect structure (random intercept for subject, Supplementary Table 10). The simplified model revealed a positive main effect of SMA theta frequency (β = 0.26, SE = 0.09, p_FDR_ = .015) in high synchronizers. Furthermore, a significant interaction effect between IFG rate ✕ STG-IFG coupling (β = −0.24, SE = 0.08, p_FDR_ = .015) indicated that IFG rate predicted speech tracking more strongly in individuals with high STG-IFG coupling. Specifically, lower IFG rates were related to stronger tracking. Additionally, we observed an interaction effect of the endogenous theta frequencies in SMA and STG (β = - 0.23, SE = 0.09, p_FDR_ = .046). For individuals with low endogenous theta frequency in SMA, speech tracking was unaffected by the theta frequency in STG. However, for individuals with higher theta frequency in SMA, speech tracking varied with theta frequency in STG such that tracking was highest in individuals with lower STG theta frequencies.

In summary, the comparison between HG and STG models reveals that the endogenous theta frequency of STG had a direct association with speech tracking, while the endogenous theta frequency of HG only played a role by interacting with other variables (such as endogenous motor frequencies and coupling). The comparison of low and high synchronizers suggests that high synchronizers exhibit stronger rhythmic speech motor system engagement during tracking (i.e. effects of endogenous motor frequencies and STG-IFG coupling).

### Similar patterns in behavioral and neural measures

Both behavioral and neural analyses exhibited similar patterns (Figures 2B and 6), with effects of the motor system and auditory-motor coupling observed predominantly in individuals displaying stronger behavioral auditory-motor synchronization (high synchronizers). Despite similar patterns of results for behavioral and neural measures, no correlations were observed. We computed partial correlations between speech comprehension and speech tracking, while controlling for syllabic rate which correlates with comprehension and tracking. The partial correlations revealed no significant effects (HG: rho = −0.03, p = .679, STG: rho = −0.06, p = .358). Regarding preferred rates, we hypothesized that the behavioral measures of the auditory and motor rates and auditory-motor synchronization reflect behavioral readouts of the underlying – supposedly – oscillatory properties of the corresponding brain systems. However, correlation analyses showed no significant relations between the corresponding neural and behavioral rates (all ps > .05).

## Discussion

Preferred rates of an individual’s endogenous (resting-state) auditory and motor brain rhythm predict their speech tracking in auditory cortex during continuous listening. This is our main result, and supports a framework of endogenous oscillations shaping perception. It refines models of auditory speech tracking and auditory-motor interactions, revealing that distinct processes may be recruited to different degrees for different parts of the population. Specifically, we find that across the population, speech tracking in higher-order auditory cortex (STG) was predicted by individual rates of endogenous theta brain rhythms in this area. Furthermore, individual rates of endogenous theta brain rhythms of speech motor cortices (particularly SMA, and IFG through interactions with STG-IFG coupling) predicted speech tracking in auditory association areas−but only in individuals with behaviorally quantified high auditory-motor synchronization.

A slightly different picture emerged for primary auditory cortex (HG). No main effect of the rate of the endogenous HG rhythm was observed, neither across the population nor separately in low or high synchronizers. In high synchronizers, as obtained for auditory association cortex, endogenous rates of theta brain rhythms of speech motor cortices (particularly IFG) predicted speech tracking in HG. Primary auditory cortex thus seems more receptive to external stimulation rates and less rhythmically constrained by its endogenous theta brain rhythm than STG. Together with our behavioral findings (which paralleled the neural results), the data suggest that during speech comprehension, individuals with higher auditory motor synchronization use motor top-down predictions through coupled auditory-motor oscillators. In contrast, low auditory-motor synchronizers rely more on auditory oscillatory populations and additionally more heavily recruit working memory processes.

### Speech comprehension and tracking decline at higher syllabic rates

As expected, we found that comprehension decreased at higher syllabic rates (above 11 syllables/sec). Although this decline was observed at slightly higher rates than previously reported in studies using simpler stimuli (e.g. words or short sentences; ,12,24,52), recent studies have observed similar effects (29,53,54). The shallower decline in comprehension performance across rates for more complex speech stimuli may be explained by enhanced comprehension due to linguistic predictability particularly at faster syllabic rates (29). Note that auditory cortex showed amplitude envelope tracking even at non-intelligible rates, and unlike Ahissar et al. (24) we observed no correlation of comprehension and tracking (after controlling for confounders like stimulus difficulty, i.e. sentence length, compression). However, our findings are in line with previous studies pointing towards a more complex link between comprehension and tracking (53,55–57). Furthermore, we acknowledge potential limitations due to a lack of trial-wise neural data analysis reducing statistical power.

### The rate of endogenous brain rhythms in STG predicts speech tracking

Endogenous theta rhythms in auditory cortex are believed to phase-align to the acoustic envelope of incoming speech (11,14,15,31,58). Our study, to our knowledge for the first time, establishes a direct relationship between an individual’s rate of the endogenous theta rhythm (measured during rest) and speech tracking in auditory association cortex (STG) during comprehension. We found that lower endogenous theta frequencies in STG were consistently associated with stronger speech tracking. This was shown in both high and low synchronizers in the general model. However, effects were stronger in low synchronizers, as seen in the group models. This finding directly suggests recruitment of an oscillatory population during speech tracking (14,32,34,37). In contrast, the endogenous theta frequency of primary auditory cortex (HG) was not predictive of its speech tracking. We speculate that due to a stronger preference for speech signals in STG (30,59), the role of its endogenous rhythm may be more relevant to speech tracking. Intracranial results have demonstrated that parts of STG preferentially represent “syllable-level temporal structure” (59), whereas HG reflects less complex acoustic features (59,60). Furthermore, oscillatory neuronal populations in STG may be involved in speech segmentation (14,30), with possibly weaker oscillatory properties in HG.

### Endogenous motor cortex rhythms predict speech tracking in high synchronizers

Endogenous speech motor cortex theta rhythms reliably predicted speech tracking in high, but not in low auditory-motor synchronizers. (Note that we define SMA and IFG as speech motor areas in this study.) The endogenous theta frequencies of SMA and IFG exhibited contrasting effects on speech tracking: higher SMA rates were associated with higher tracking, higher IFG rates with lower tracking. This is consistent across tracking in auditory association and primary auditory cortex (STG and HG; Fig 5 and 6). Interestingly, STG particularly showed effects of the rates of SMA (less so for IFG), while the opposite was observed for HG. Given SMA’s role in temporal processing (4,61–63) and IFG’s involvement in temporal sequencing and speech motor processing (4,64), these findings indicate that high synchronizers may rely more heavily on temporal motor predictions for speech processing, compared to low synchronizers. This interpretation is further supported by our behavioral observation of increased comprehension performance in high synchronizers. The endogenous theta frequency in IFG showed a negative effect, indicating enhanced tracking with lower endogenous rates. Given the proposed direct connection between IFG and STG (4), observing the same direction of effects may imply a synergistic effect of (the endogenous frequencies in) these two areas on tracking. In contrast, the positive effect of the endogenous theta frequency in SMA on tracking paralleled our behavioral findings and showed the expected direction of the effect. The discrepancies between the SMA and IFG effects may reflect differences in network connections, as well as whether the areas are connected in an excitatory or inhibitory manner (65). Further research is necessary to elucidate the distinct relation of endogenous IFG and SMA rhythms to speech tracking. The finding of selective motor effects for high synchronizers is in line with previous work (27,29,39,44,66,67). Particularly, Assaneo, Rimmele et al. (39) proposed, based on behavioral data and a neural computational model, that oscillatory auditory-motor coupling was engaged preferentially in high synchronizers. Here, in low synchronizers, we only detected an interaction effect between the endogenous theta peak frequencies in IFG and HG. High compared to low synchronizers may not only rely more strongly on the recruitment of the speech motor system but also more extensively engage additional areas like (parts of) the SMA, reflecting other aspects of temporal processing or alternative processing routes (4).

### Auditory-motor coupling interacts to affect speech tracking

Overall, we observed similar patterns for theta phase auditory-motor coupling between different motor (IFG, SMA) and auditory (HG, STG) areas (Fig S2), with significant auditory-motor coupling at all syllabic rates. Coupling, however, decreased from 5Hz to 11Hz and slightly increased again at faster rates. The findings align with reports of a “sweet-spot” for auditory-motor coupling around 4.5 Hz (17). Furthermore, such a sensitivity for certain frequencies in the theta- and beta-range has been previously shown for the auditory cortex (68) and for the motor cortices (as reflected in their endogenous rhythms; ,34,36). Speech tracking was predicted by coupling only in high synchronizers, who showed an interaction effect of IFG-STG coupling and the IFG rate on STG tracking. Specifically, effects of the endogenous IFG theta rate on speech tracking were more pronounced in individuals with high auditory-motor coupling strength. Our results are limited by the focus on phase-phase theta coupling. Further research is required to understand oscillatory processing routes within a spectrally and spatially more complex auditory-motor network.

### Behavioral findings parallel the neurophysiological results

Replicating our two behavioral experiment(s) (29), we show that higher speech comprehension was related to higher individual spontaneous motor production rates, auditory-motor synchronization, and preferred auditory rates (in our previous study, the later was a trend that did not survive multiple-comparison control). A 3-way interaction between these variables suggests a complex interplay of the behaviorally assessed preferred rates and auditory-motor synchronization. Interestingly, the behavioral findings remarkably paralleled our neuronal results. In low synchronizers, comprehension was predicted by the preferred auditory rate and, interestingly, by working memory scores. In contrast, in high synchronizers, comprehension was predicted by the preferred auditory rate, and, crucially, the motor production rate and the interaction of auditory and motor rates.

Despite similar effects in behavior and the brain, we observed no correlations between the corresponding measures. Speech comprehension is intricate and shaped by numerous interacting variables. While the neuronal dynamics underlying speech tracking are crucial, they likely represent just one aspect of the number of computations—including linguistic and situational predictions, as well as working memory—that facilitate speech comprehension.

## Conclusion

In a large sample of participants, we demonstrate that *endogenous* theta rhythms of STG predict speech tracking. Interestingly, endogenous theta rhythms of speech motor areas (SMA, IFG) were predictive of speech tracking only in those individuals showing high behavioral auditory-motor synchronization. Our findings are consistent with an oscillatory model of auditory-motor interactions during speech comprehension. While some individuals recruit the speech motor system (SMA, IFG), likely providing temporal predictions to enhance comprehension, others predominantly rely on auditory processing, possibly recruiting working memory processes, more strongly. Furthermore, our findings suggest differences of primary auditory and association areas in their oscillatory characteristics relevant for speech tracking and in their processing routes.

## Materials and Methods

Our magnetoencephalography (MEG) and behavioral experiment entailed three separate sessions: a behavioral session, an MEG session with behavior, and a structural magnetic resonance imaging (MRI) session. The study was approved by the local ethics committee of the University Hospital of the Goethe-University Frankfurt (number: 2021-509) in accordance with the Declaration of Helsinki.

### Participants

Overall, the data of N=57 participants were analyzed (age: M = 26.9, SD = 5.4; 32 female, self-report of gender, i.e. German “Geschlecht”). The initial sample consisted of N=60 participants (N=3 participants were excluded because of technical issues during recording, N=2, and because of an average performance of 3 standard deviations below average in the baseline condition (5 syllables/s) of the speech comprehension task, N=1). As assessed by self-report, participants had no history of neurological or psychiatric diseases and had normal hearing, as well as normal (or corrected-to-normal) vision. All participants were native speakers of German and right-handed. Prior to each session, participants provided written informed consent. At the end of the final session, participants received monetary compensation. Due to analysis specific exclusion criteria, different numbers of subjects were included in the different analyses. Specifically, participant numbers were affected by (1) the SSS test and (2) the extraction of the individual theta peak frequency at auditory and audio-motor brain areas. The number of participants included in each analysis and the exclusion criteria are stated in the respective analysis methods and results section – if they diverge from the general sample of 57 subjects.

### Stimuli

In two experimental tasks (speech comprehension task and preferred rate task) participants listened to naturalistic sentences. The sentences were sourced from German books (N_sentences = 306; 8 talkers; source: zeno.org) and audiobooks (N_sentences = 138; 3 talkers; source: Librivox.org), respectively. Sentences from books were recorded by three native talkers of German at the MPIEA in a sound-attenuated booth using MatLab R2017a on a Windows 7 Pro (64-bit) and a Neumann U87i studio microphone, and A/D conversion at 44.1 kHz. Talkers delivered the sentences at their normal, slowest, and maximal speaking rate, while prioritizing proper articulation over speed. All sound files, including audio books, underwent processing using Praat (6.0.40). Long pauses (>300ms) were removed to prevent inaccurate rate (syllables/second) estimates.

Three stimulus lists were generated from the sentence materials for both the speech comprehension task (300 sentences) and the auditory rate preference task (132 sentences). Time compression or expansion was applied to all sentences to create different syllabic rate conditions. Sentences were randomly selected (without replacement) from the total pool of sentences based on two main criteria: the degree of compression required to obtain the desired syllabic rate (compression factor < 3) and the duration of the sentence after compression (> 1s; only applied to comprehension task). No sentence repetitions occurred within each stimulus set across tasks. Time compression or expansion was performed using the Pitch Synchronous Overlap and Add (PSOLA) algorithm in Praat, and all stimuli were standardized to a root mean square (RMS) amplitude at 69 dB (see Fig 3).

## Experimental tasks

### Intelligibility task

To measure speech comprehension, participants performed an intelligibility task (Fig 4A). On each trial, participants listened to a sentence via headphones and verbally repeated it as accurately as possible. All responses were recorded, and participants stopped the recording via button press (left or right index finger), initiating the interstimulus interval.

Sentences were presented at five syllabic rates (5.00, 11.00, 14.00, 16.00, 17.50 syllables/s) with 60 different sentences each, totaling 300 trials. Trials with different syllabic rates were pseudorandomized within blocks of 30 trials, with self-paced breaks between blocks.

### Spontaneous speech synchronization (SSS) test

To assess auditory-motor synchronization, participants performed the spontaneous speech synchronization (SSS) test (Fig 1A; for detailed description see the General Introduction, study 2 and Assaneo et al., 2019). In two trials, participants continuously whispered a syllable (/te) for 80 seconds and aimed to synchronize their own motor output to a stream of syllables. Their whispering was recorded. The auditory stimulus progressively increased in rate from 4.3 to 4.7 syllables/s (increments of 0.1 syllables/s) every 60 syllables. Participants’ syllable production was masked by the simultaneously presented auditory syllable train.

### Speech production task

The individual spontaneous speech motor production rate was measured by asking participants to freely speak “as they would naturally” (Fig 1B). Participants were prompted with 6 questions/statements to facilitate continuous speech production (6 trials; own life, preferences, people, culture/traditions, society/politics, general knowledge; see Supplementary Table 2; (69). Participants read the question/statement and initiated the speaking period (30 s) via button-press. White noise was presented via headphones to minimize auditory feedback. Breaks in between trials were self-paced.

### Auditory rate preference task

The preferred auditory rate was assessed using a two-interval forced choice (2IFC) task (Fig 1C). On each trial, we presented two versions of the same sentence, randomly ordered, differing only in syllabic rate. After stimulus presentation, participants indicated which of the two stimuli they preferred via button press. Stimuli were presented at twelve syllabic rates from 3.0 to 8.5 syllables/s (in steps of 0.5 syllables/s).

### Digit span test

Working memory capacity was quantified using the forward and backward (70) digit span test. Participants listened to digit spans and typed in their responses after listening (71). The test comprised seven levels, ranging from two/three to nine digits with two items at each level. The procedure started with the shortest digit spans (forward: three digits, backward: two digits) and stopped as soon as participants failed to repeat both spans of the same length or when the two longest digit spans were reached.

### Procedure – Behavioral session

Participants were seated in a sound-attenuated experimental booth equipped with a Fujitsu CELSIUS M740B PC. Stimuli were presented using Psychtoolbox (Brainard, 1997) in Matlab (9.7.0.1471314, R2019b) and insert earplugs (ER3C Tubal Insert Earphones; Etymotic Research). Participants’ speech and whisper was recorded using a gooseneck microphone (MX418 microflex gooseneck microphone, Shure) and behavioral responses were collected using a standard keyboard. Stimuli were presented at ∼70 dB in the preferred auditory rate and digit span tasks. For the spontaneous speech production task and the SSS-test, loudness could be adjusted using a volume control knob on the sound card. Participants were instructed to increase the volume until their own speech or whisper became inaudible, while still being comfortable. The rationale behind this procedure is to isolate motor production by suppressing auditory feedback.

All participants started with the spontaneous speech motor production rate task to avoid priming effects. Next, participants performed either the SSS-test or the preferred auditory rate preference task, the order was randomized across participants. Finally, all participants finished with the digit span task, followed by the questionnaire (demographics and musicality).

### Procedure – MEG session

After application of EOG and ECG channels, participants were seated in the MEG. The stimuli were delivered binaurally using insert earplugs (EARTONE Gold 3A insert earphones; Ulrich Keller Medizin-Technik) and the Matlab (R2017a) software with the Psychtoolbox (3.0.14; Brainard, 1997) extension on a Fujitsu-Technology CELSIUS R940power PC. Participants’ responses were recorded using a button box (Current Designs Package 932). Fiducials were attached to participants’ nasion and the preauricular points to continuously measure head position.

First, participants performed the intelligibility task. The task was split into 10 blocks (∼5 minutes each). Second, participants completed an auditory localizer task in which they passively listened to a sequence of sounds (pure tones: 0.4 s tone duration; 250 Hz and 1000 Hz, 100 repetitions, jittered intertrial interval 0.5–1.5 s; ∼5 min). Third, participants performed a motor localizer task (∼12 min) during which they repeatedly articulated syllables without vocalizing. On each trial, one out of three syllables (/pa/, /ta/, /sa/) was articulated for 3 s and each syllable was repeated in 50 trials. Finally, we recorded resting state activity (∼5 min 30 sec). Throughout the experiment, participants were instructed to hold their gaze at a fixation cross (except during instructions). In total, the MEG session had a duration of roughly 180 minutes, consisting of 120 minutes recording time and 60 minutes of preparation time and scanning pauses.

MEG data were acquired at a sampling rate of 1200 Hz using a 275-channel whole-head MEG machine (Omega 2005, CTF Systems Inc.) in a magnetically shielded room. During scanning, online denoising (higher-order gradiometer balancing) and online low-pass filtering (cut-off: 300 Hz) were applied. Furthermore, participants’ head position was measured continuously using fiducials and the Fieldtrip toolbox (72, version 20220617), allowing for position adjustment between blocks as well as continuous head movement correction at the analysis stage.

### Procedure – MRI session

Individual structural MRI scans (standard 1 mm T1-weighted MPRAGE) were obtained using a 3 Tesla scanner (2 participants were scanned on a Siemens Magnetom Trio scanner; all other participants were scanned on a Siemens Magnetom Prisma, scanner Siemens, Erlangen, Germany. Vitamin E capsules were used to mark anatomical landmarks (nasion, left and right pre-auricular points) to align MRI and MEG data for source reconstruction.

## Behavioral Analysis

### Speech comprehension

Participant’s responses were transcribed manually. To quantify comprehension accuracy, we compared the responses to the original sentences using a sequence matcher algorithm (Python built-in sequence matcher), which quantifies the similarity between two sequences by (order of) item, i.e. letter. The output of the sequence matcher was a percentage for each sentence.

### Preferred auditory rate

From the 2IFC task, we derived the preferred frequencies for each trial. This distribution of preferred frequencies across trials was fitted using a Gaussian function. From the fitted Gaussian, the peak parameter was extracted and defined as individual preferred auditory rate.

### Spontaneous speech motor production rate

The spontaneous speech motor production rate was quantified as articulation rate (i.e. number of produced syllables divided by trial duration, excluding pauses > 300ms). To segment the continuous speech recordings into syllables, syllable nuclei were detected automatically using Praat (73). The spontaneous speech motor production rate was computed for each trial separately and then averaged across trials.

### Auditory-motor synchronization

The data from the SSS-test were analyzed according to the protocol of Lizcano-Cortés et al. (50). First, data quality was ensured by confirming that participants whispered (i.e. no vocal cord activation) and that audio files were not corrupted (i.e. noise interference). For the analysis, we applied the scripts provided by the authors (50). Auditory-motor synchronization was measured as the phase-locking value (PLV) between the speech envelope of the produced motor signal and the cochlear envelope of the syllable stimulus. For the syllable train, the cochlear envelope was extracted using the Chimera toolbox (auditory channels: 180–7,246 Hz) (74). For the produced motor signal, the amplitude envelope was estimated using the Hilbert transform (hilbert.m function in Matlab). Both envelopes were down-sampled to 100Hz and bandpass filtered between 3.5 and 5.5 Hz. From the filtered signal, we extracted the phase (using hilbert.m and phase.m functions in Matlab), computed the PLV for time windows of 5s (overlap 2s) and averaged across time windows. This procedure was completed for both experimental runs separately. To test for consistency between runs, a linear regression was fitted to the data (independent variable: PLV of run 1, dependent variable: PLV of run 2). Participants with PLV pairs outside the 95^th^ confidence interval were excluded from further analysis. For the remaining participants, PLVs from both runs were averaged. The mean PLVs were subjected to a Gaussian mixture model to partition participants into high and low synchronizers. While we obtained PLVs for all N = 57 participants, ten could not be classified clearly as high synchronizers (probability being a high synchronizer = 45-55%). Accordingly, the final sample contained N = 27 high synchronizers and N = 20 low synchronizers.

## MEG analysis – speech comprehension task

### Speech envelope extraction

For the speech signals presented during the comprehension task in the MEG, we extracted the cochlear envelope using the Chimera toolbox (auditory channels: 180–7,246 Hz) (74). Specifically, the spectrograms were computed for multiple frequency bands (auditory channels: 180–7246 Hz) and the absolute values then averaged to obtain the broadband speech envelope. Broadband envelopes were down-sampled to 100 Hz to match MEG signals.

### Preprocessing

During MEG preprocessing, the data were bandpass filtered (0.5–160 Hz, Butterworth filter; filter order 4) and line-noise was removed (49.5-50.5, 99.5-100.5, 149.5-150.5 Hz, two-pass; filter order 4). We applied a semi-automatic artifact rejection procedure to detect jump, muscle, and threshold artifacts. For jump and muscle artifacts, data were filtered to optimize artifact detection (muscle: 110-140 Hz, jump: median filter) and then z-transformed per sensor and time point. Trials were rejected if they surpassed a priori defined thresholds (jump: z=45, muscle: z=15). For the threshold artifacts, trials were rejected if the range (minmax difference) of activity at any channel surpassed an a-priori defined threshold (threshold = 0.75e−5). The data were down-sampled to 500 Hz and epoched (−1000ms to +100ms after trial offset), resulting in epochs of variable length. Using the continuous fiducial measures, trials containing head movements larger than 4 mm were identified and rejected. Data from the separate recording blocks were concatenated into one large file and sensors with high z-values (z > 2) were rejected. Finally, eye blink, eye movement and heartbeat artifacts were corrected using independent component analysis (infomax algorithm; Makeig et al., 1996). Data were further down-sampled to 100 Hz for computational efficiency.

### Source localization

Using the three anatomical landmarks (nasion, left, and right preauricular points), individual T1-weighted MRI images were co-registered to the MEG coordinate system in a semiautomatic procedure. T1-weighted MRIs were segmented into three tissues (white matter, grey matter, cerebrospinal fluid) to construct single-shell volume conduction models (headmodel) (75) and warped into MNI space to compute individual grids by inverse warping the template grid (grid resolution: 5mm) onto individual anatomical scans. Using the individual grids and volume conduction models, we computed individual forward models to reconstruct source activity.

For source reconstruction, we used a Linearly Constrained Minimum Variance (LCMV) Beam-former (array-unit LCMV) (76). Firstly, we computed the covariance matrix across all trials and created a common filter for all conditions using the individual forward models. The lambda regularization parameter was set to 1% and time-series were extracted for all three dipole orientations per voxel. Secondly, trial data was source-localized by projecting it through (i.e. multiplying it with) the common filter for each condition separately. Computing the common filter across all conditions -instead of separate filters for each condition-ensures that differences between conditions reflect differences in source activity, instead of differences in the spatial filters itself.

### Regions of interest

For the speech tracking analysis, regions of interest (ROI, as defined by the Automated Anatomical Labeling (AAL) atlas (77)) included primary and non-primary auditory cortex, comprising Heschl’s gyrus (HG) and superior temporal gyrus (STG), respectively. Both areas are crucial for speech perception, with the STG showing increasing responsiveness to complex acoustic speech features compared to HG, as supported by neurocognitive models and empirical studies (59,78–80). Notably, speech tracking has been consistently observed in both HG and STG across multiple studies (7,14,81–85).

For auditory-motor coupling, in addition to HG and STG, the ROIs also included two speech motor areas that are well-established in the speech motor network. Inferior frontal gyrus (IFG), a key component of the core language network (79) which might contribute to the auditory-motor coupling during speech perception (6,44), exhibits direct structural connections with the auditory cortex (4,86) and is involved in various speech processing subroutines such as phonological encoding (87) or semantic and sentential information integration (88). Importantly, IFG synchronizes to speech envelopes (44,82) and shows differing degrees of synchronization depending on individual auditory-motor speech coupling (44). Interestingly, the auditory-motor coupling of IFG has been proposed to reflect neural oscillatory activity (89). The supplementary motor area (SMA), although not traditionally considered a language area, plays a central role in speech motor control (1) and, more recently, has been recognized to contribute to higher-level processes during speech processing (4). Together with the cerebellum and basal ganglia, SMA processes temporal structure (90) and sensory temporal information more generally (61–63). Notably, SMA is particularly implicated in adverse listening situations (91), possibly through top-down mechanisms, predicting upcoming sensory events (63,92).

We extracted the corresponding parcels for all ROIs (13+14: IFG tri.; 19+20: SMA, 79+80: HG, 81+82: STG) from the source-localized data using the AAL atlas. To further reduce the dimensionality of the data, principal components were computed across voxels within each parcel. To this end, for each trial, all voxel time-series within a parcel were stacked (three time-series, i.e. for each dipole orientation, per voxel) and principal components and their explained variance were extracted. Based on previous work (93), the further analysis only included the first three principal components.

### Mutual information – speech tracking

Speech tracking was estimated using information theory (82,94,95). Specifically, we computed gaussian-copula mutual information (GCMI, 51) to assess statistical dependencies between speech envelopes and source-localized neural activity in auditory and motor ROIs. GCMI capitalizes on the concept of gaussian copulas which is a statistical description of the relationship of two variables regardless of their respective marginal distributions. Mutual information (MI) can be computed as the negative entropy of a statistical copula. This facilitates estimation of MI because 1) no assumptions of the marginal distributions of the variables are needed and 2) the computation is efficient as a Gaussian parametric estimation can be employed. A further advantage of the GCMI approach is that it lends itself naturally to dealing with multivariate data (51). This allowed us to use phase information of speech envelopes and source-localized activity for multiple frequency bands.

For the estimation of GCMI, we applied a continuous wavelet transform for 28 frequencies (3.5 to 20 Hz) to the source-localized principal components for each trial using *cwtfilterbank.m* in Matlab. To avoid onset effects, filtered principal components were epoched again from 200ms relative to stimulus onset until trial end. From the complex signals, we first extracted the phase information (real and imaginary parts) and then copula-normalized the parts separately. Finally, the normalized components were concatenated within each condition. This yielded two frequency-by-samples vectors per condition, i.e. source time-series. Importantly, the same procedure (continuous wavelet transform, epoching, extraction of phase information, concatenation) was applied to the speech envelope data with the only difference that each trial only consisted of one time-series per trial (no PCA). Analogous to the source time-series, this resulted in two speech time-series per condition (real and imaginary parts).

In a multivariate analysis, we estimated GCMI between 2 speech time-series and 6 source time-series for each parcel and frequency. To account for stimulus-brain lags, we applied our GCMI estimation at various positive delays (0ms to 300ms, 10ms steps).

### Mutual information – Auditory-motor coupling

To examine the hypothesis that audio-motor coupling strength predicts speech tracking (and comprehension), we estimated audio-motor coupling using GCMI. This analysis was identical to the speech tracking analysis above with respect to frequency analysis and GCMI parameters. The important difference lies in the selection of time series: here GCMI was computed between MEG time courses from two different brain areas. Specifically, we computed GCMI between HG or STG (both left and right) and IFG pars triangularis or SMA (both left and right).

We tested whether auditory-motor coupling was higher than chance-level within each condition by comparing the true MI values against the 99^th^ percentile of surrogate data. To create surrogate data, we segmented the concatenated trials of each condition into segments, shuffled the segments for each (N=500) iteration and estimated GCMI between source time-series and shuffled speech time-series. Importantly, segment length differed between conditions and was set at 80% of the cycle length of the stimulation frequency of each condition (i.e. 5Hz = 160.0ms, 11Hz = 72.72ms, 14Hz = 57.1ms, 16Hz = 50ms, 17.5Hz = 45.7ms). We used the surrogate distribution to create z-transformed GCMI values, thus subtracting true GCMI values by the mean surrogate distribution and dividing the result by the surrogate’s standard deviation. The normalization was performed within participants for each parcel, condition, and frequency. GCMI values varied systematically as a function of delay and condition; we extracted GCMI for each condition at the delay that maximized the GCMI values.

## MEG analysis – resting state

### Preprocessing

For the resting state data, preprocessing was similar to the main task but adjusted in a few aspects to match the preprocessing procedure employed in previous studies (36). Specifically, all parameters for artifact rejection were identical to the main task of the current study. The data were detrended, down-sampled to 250 Hz and epoched into 0.8 s trials, which resulted in an average of M=380 trials (sd=59.8, min=121, max=412). Sensors were rejected based on the neighborhood-ratio using the *hcp_qc_neighstddratio.m* function (Human Connectome Project, WU-Minn Consortium), which is defined by the sensors noise level relative to the noise level of neighboring sensors (Sensor SD—NeighborSD/NeighborSD). Sensors were rejected if the ratio exceeded a value of 0.5. For trial rejection based on head movements and the independent component analysis, the same procedures as in the main data of the current study were implemented.

### Common filter for source projection

We used the same headmodels as in the main analysis. Forward models were obtained using the same procedure as for the main task, but with a different grid (resolution = 10mm; see study 1 and (34)). For source projection, we used an LCMV Beamformer. First, we computed the covariance matrix across all trials for each individual. Second, the common filter was generated using the previously computed forward models.

The lambda regularization parameter was set to 7% and the optimal dipole orientation for each voxel was extracted (single value decomposition). The beamformer coefficients were obtained in preparation for the later source projection of Fourier spectra.

### Spectral analysis and source projection

Complex Fourier spectra were computed in sensor space for individual epochs (0.8s) at 1-120 Hz (logarithmic frequency resolution) and each participant. Each epoch was zero-padded to a duration of 2s before applying a multitaper spectral analysis (DPSS multitaper, 2 tapers, +/- 2 Hz spectral smoothing). The complex spectra were projected into source space by multiplication with the previously obtained LCMV coefficients, summing across tapers for each trial. Source-localized spectra were ratio-normalized (the spectral power for each voxel and segment was divided by the average power across all voxels and segments) and subtracted by 1, yielding values above/below zero. To reduce dimensionality, we parcellated the data into 115 parcels using the AAL atlas by averaging power spectra across voxels within each parcel.

### Clustering spectral data

For each participant and parcel the source-localized single-trial Fourier spectra were subjected to the k-means algorithm. The k-means algorithm treated each trial’s spectrum as a 42-dimensional space and grouped all trials into *k* mutually exclusive spectra according to spectral (dis)similarities. These spectral clusters represent the spectral modes the brain area engages in over time. We set *k* to 10 (see: 34), used the Cosine distance metric, and initiated the k-means algorithm 10 times (max. 100 iterations). To quantify the proportion of trial-spectra –as determined by the k-means algorithm– belonging to each cluster, k-means clusters were subjected to a Gaussian Mixture (GM) Model algorithm.

Prior to the group-level analysis, we determined the optimal number of clusters across participants for each parcel. We evaluated the Silhouette criterion (1000 iterations) for solutions between 1-15 clusters and selected the number of clusters characterized by the highest Silhouette value. To obtain group-level spectra for each parcel, single-subject clusters were stacked and then fed into the k-means algorithm. The results were again subjected to the GM Model algorithm.

### Selection of individual source-localized theta peaks

We extracted individual peak frequencies from the theta clusters for HG, STG, SMA, and IFG (see above for motivation of ROIs). Spectral clusters were defined as theta cluster if their peak frequency was within the range of 4 to 8 Hz. To this end, first, we extracted single-subject spectral clusters that contributed to the theta group-level cluster in question. From the single-subject clusters, we computed the peak frequency (henceforth *endogenous theta frequency of area X*). If multiple clusters from one participant contributed to the group cluster, the cluster with the highest peak amplitude was selected. On the group-level, HG was characterized by two theta clusters. We extracted the individual clusters contributing to the lower group cluster because it was contributed to by N=56 participants, compared to N=44 for the higher cluster.

## Statistical analysis

To assess the hypothesized links between speech comprehension and speech tracking and the corresponding predictor variables, we computed Generalized Linear Mixed Models (GLMM). Models were computed using R (version 4.1.3, 2022-03-10) set up in Rstudio (version 2022.2.1.461). For the GLMMs, we used the R package *lme4* (version 1.1-35.1). Tables and effects are visualized using *sjPlot* (version 2.8.15) and *ggplot2* (version 3.4.4). For all models, continuous predictor variables were z-transformed. All results are corrected for multiple comparisons using false discovery rate (FDR) correction, unless stated otherwise.

First, for the behavioral GLMM, trial-wise speech comprehension performance (% words correct) was regressed against syllabic rate, preferred auditory rate, spontaneous speech motor production rate, PLV, working memory score, compression factor, sentence length (in syllables), and stimulus order. We were particularly interested in the interaction effect of PLV ✕ preferred auditory rate ✕ spontaneous speech motor production rate. Thus, in addition to main effects, the model contained a three-way interaction. As random effects, we introduced a by-participant random slope for syllabic rate and a random intercept for trial-ID. Overall, the model (N = 57) explained 47.8% of the variance. To reduce the complexity, we computed the model separately for high and low synchronizers, which allowed us to reduce the three-way interaction to a two-way interaction (no PLV term). These high (N = 27) and low (N = 20) synchronizer models explained 47.1% and 48.1% of the variance, respectively.

Second, we designed GLMMs that computed the relationship between speech tracking and the neural predictor variables. All neural variables (speech tracking, endogenous rates, auditory-motor coupling) were averaged across hemispheres. We computed two separate models, one focused on HG (“HG general model”) and one on STG (“STG general model”). Participants were excluded from this analysis if they did not display single-subject spectral peaks in the respective brain areas. In the HG general model (N = 54), speech tracking in HG was regressed against syllabic rate, auditory-motor coupling (IFG-to-HG and SMA-to-HG) and the endogenous theta frequencies of HG, IFG, and SMA. The model included two three-way interactions between auditory-motor coupling and the corresponding endogenous theta frequencies. The STG general model (N = 50) was identical except that all HG variables were exchanged for STG variables (endogenous rate and coupling). Overall, the HG model explained 24.8% of the variance, the STG model 35.6%. Analog to the behavioral GLMM, additionally, we first added the PLV as predictor to the general models for HG and STG and then computed the models separately for high and low synchronizers. Partitioning the sample into high and low synchronizer groups resulted in the following sub-samples: HG_highs: N = 27, HG_lows: N = 20, STG_highs: N = 27, STG_lows: N = 19.

Third, partial correlation analyses were computed to quantify the link between speech comprehension and speech tracking, while controlling for the effect of syllabic rate. The correlation analyses were computed for speech tracking in STG and HG.

## Acknowledgements

We thank Dr. Klaus Frieler for valuable advice on the statistical analysis, and Hong Ngoc Tran Thi, Daniela van Hinsberg, and Aayush Marishi for help with data collection.

