## Supplementary Material for "Endogenous auditory and motor brain rhythms predict individual speech tracking"

**Supplementary materials**

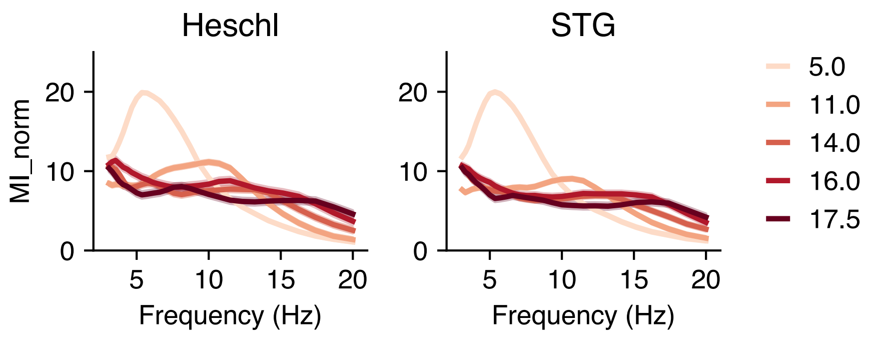


**Supplementary Figure 1. Normalized MI spectra for speech tracking in auditory cortex.** Line graph illustrates normalized MI spectra in Heschl's gyrus (HG, left) and posterior superior temporal gyrus (STG, right). For both panels, lines are color-coded according to the syllabic rate of sentence stimuli and shaded error bars represent standard error of the mean across participants.

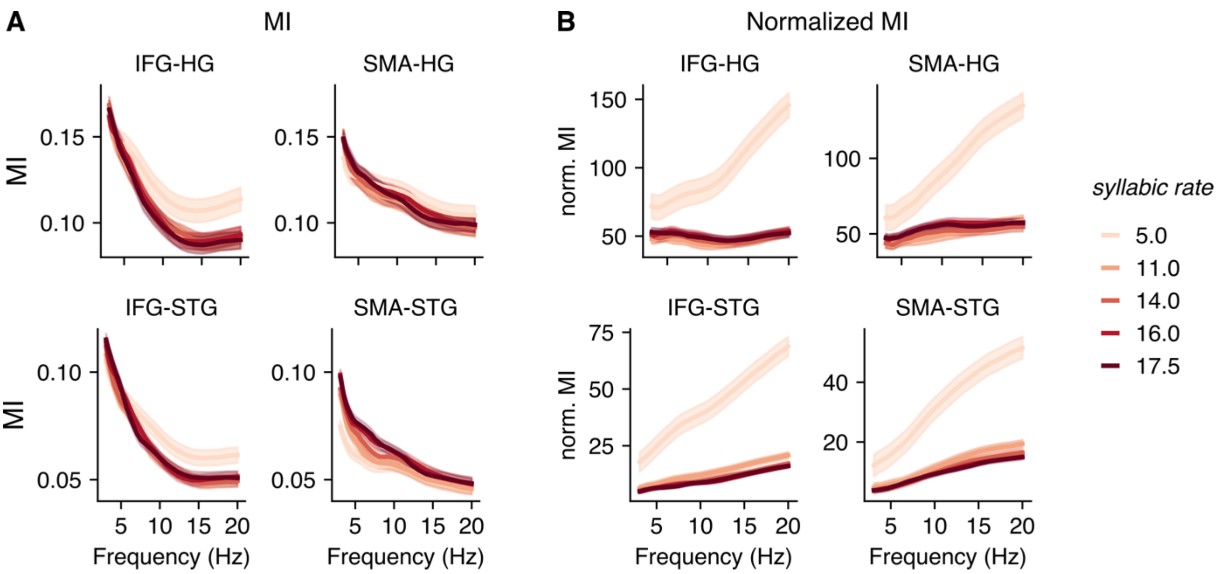


**Supplementary Figure 2. Raw MI spectra for auditory-motor coupling.** Line plots illustrate raw (**A**) and normalized (**B**) auditory-motor coupling (MI) between different auditory and motor brain areas as a function of frequency: IFG-HG, SMA-HG (top), IFG-STG , SMA-STG (bottom). Lines are color-coded according to the syllabic rate of sentence stimuli and shaded error bars represent standard error of the mean across participants. HG = Heschl's gyrus, STG = superior temporal gyrus; IFG = interior frontal gyrus; SMA = supplementary motor area.

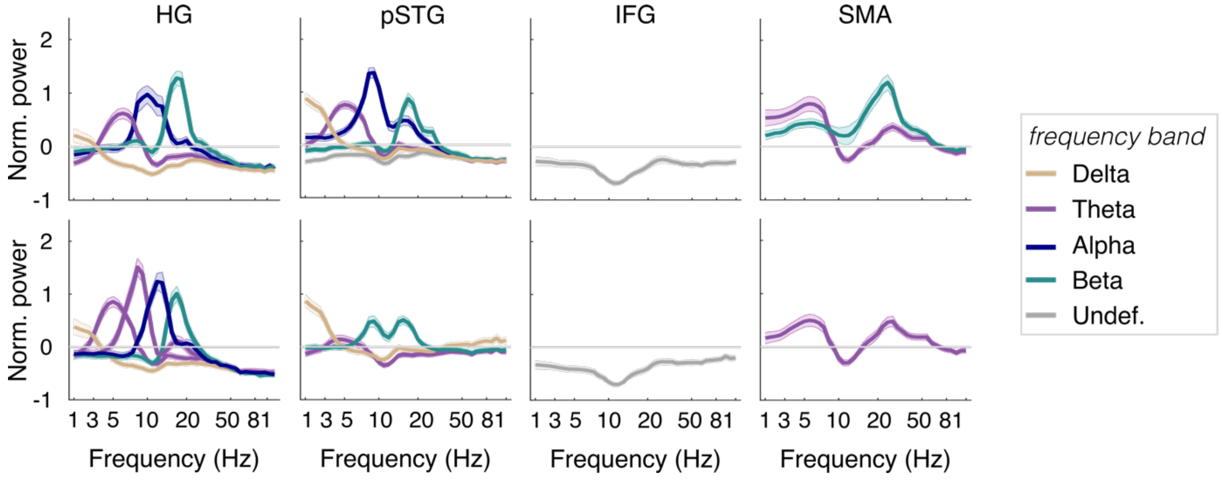


**Supplementary Figure 3. Group-level spectral fingerprints in the left (top) and right (bottom) hemisphere.** Clusters are color-coded according to their peak frequency (legend on the right). Shaded error bars reflect standard error of the mean across participants. HG = Heschl's gyrus; STG = posterior superior temporal gyrus; IFG = interior frontal gyrus; SMA = supplementary motor area.

**Supplementary Table 1. Sentence materials: Titles and authors of (audio)books.** List of sources from which stimuli for the tasks (speech comprehension and preferred auditory rate tasks) were constructed.

| Type | Author | Title | Source |
|---|---|---|---|
| PDF | Hedwig Dohm | Sibilla Dalmar | Zeno |
| PDF | Hedwig Dohm | Schicksale einer Seele | Zeno |
| PDF | Hedwig Dohm | Christa Ruland | Zeno |
| PDF | Dora Duncker | Großstadt | Zeno |
| PDF | Bertha von Suttner | Martha's Kinder | Zeno |
| PDF | Bertha von Suttner | Die Waffen nieder | Zeno |
| PDF | Stefan Zweig | Schachnovelle | Zeno |
| PDF | Ida Boy-Ed | Fanny Förster | Zeno |
| Audiobook | Frances Hodgson Burnett | Der kleine Lord | Librivox |
| Audiobook | Ludwig Thoma | Tante Frieda | Librivox |
| Audiobook | Ludwig Thoma | Lausbubengeschichten | Librivox |

**Supplementary Table 2. Prompts for speech production task.** Thematic questions used to facilitate natural speech production. Each item is representative of a different thematic category, as introduced by Alexandrou et al.(2016). Sentences were translated to German.

| Category | Sentence/statement |
|---|---|
| Own life | Welche Hobbies haben Sie oder hatten Sie in Ihrem bisherigen Leben?<br>What kind of hobbies do you have or have had during your life? |
| Preferences | Welche Art von Urlaubstrips mögen Sie am liebsten?<br>What kinds of vacation trips do you like? |
| People | Beschreiben Sie bitte einen Künstler, Schriftsteller oder Filmdirektor. Warum finden Sie ihn/sie interessant?<br>Describe a known artist, writer, or film director. Why do you find her/him interesting? |
| Culture/traditions | Beschreiben Sie bitte ein traditionelles Weihnachtsfest.<br>Describe a traditional Christmas holiday. |
| Society/politics | Was wissen Sie über Müll- und Recyclingregelungen in Ihrem Heimatland?<br>What do you know about garbage and recycling policies in your home country? |
| General knowledge | Was wissen Sie über Skifahren und Snowboarden?<br>What do you know about skiing and snowboarding? |

**Supplementary Table 3. Predicting single-trial speech comprehension performance.**

Predicted speech comprehension

| Predictors | beta | SE | t | $p_{FDR}$ | p |
|---|---|---|---|---|---|
| syllabic rate | -70.81 | 1.78 | -39.76 | **<0.001** | **<0.001** |
| syllabic rate$^2$ | -24.36 | 0.98 | -24.82 | **<0.001** | **<0.001** |
| PLV | 0.03 | 0.02 | 2.28 | **0.032** | **0.023** |
| motor rate | 0.03 | 0.02 | 2.02 | 0.056 | **0.044** |
| auditory rate | 0.09 | 0.02 | 5.94 | **<0.001** | **<0.001** |
| working memory | 0.07 | 0.01 | 4.94 | **<0.001** | **<0.001** |
| compression | -0.00 | 0.01 | -0.19 | 0.850 | 0.850 |
| num. syllables | -0.11 | 0.02 | -4.99 | **<0.001** | **<0.001** |
| stimulus order | 0.05 | 0.00 | 10.21 | **<0.001** | **<0.001** |
| PLV * motor rate | 0.07 | 0.02 | 4.16 | **<0.001** | **<0.001** |
| PLV * auditory rate | 0.01 | 0.02 | 0.50 | 0.708 | 0.620 |
| motor rate * auditory rate | -0.06 | 0.02 | -3.29 | **0.002** | **0.001** |
| PLV * motor rate * auditory rate | -0.06 | 0.02 | -3.58 | **0.001** | **<0.001** |

Random effects

| Group | Name | Variance | STD |
|---|---|---|---|
| Trial-ID | (Intercept) | 0.203231 | 0.45081 |
| Subjects | Syllabic rate | 0.000377 | 0.01942 |
| Residual | | 0.2944557 | 0.54264 |
| N (subs) | | 47 | |
| Observations | | 14100 | |
| Marginal $R^2$ | | 0.478 | |

**Supplementary Table 4. Predicting single-trial speech comprehension performance, separately for high and low synchronizers.**

| Predicted speech comprehension | | | | | | | | | | |
|---|---|---|---|---|---|---|---|---|---|---|
| | High synchronizers | | | | | Low synchronizers | | | | |
| Predictors | beta | SE | t | $p_{FDR}$ | p | beta | SE | t | $p_{FDR}$ | p |
| syllabic rate | -53.61 | 1.69 | -31.76 | **<0.001** | **<0.001** | -45.67 | 2.02 | -22.58 | **<0.001** | **<0.001** |
| syllabic rate$^2$ | -19.73 | 0.98 | -20.15 | **<0.001** | **<0.001** | -15.82 | 0.92 | -17.14 | **<0.001** | **<0.001** |
| motor rate | 0.08 | 0.02 | 3.73 | **<0.001** | **<0.001** | -0.02 | 0.02 | -0.97 | 0.473 | 0.331 |
| auditory rate | 0.05 | 0.02 | 2.73 | **0.010** | **0.006** | 0.08 | 0.02 | 3.93 | **<0.001** | **<0.001** |
| working memory | -0.02 | 0.02 | -0.76 | 0.506 | 0.449 | 0.13 | 0.02 | 6.08 | **<0.001** | **<0.001** |
| compression | 0.01 | 0.01 | 0.50 | 0.621 | 0.621 | -0.01 | 0.01 | -0.67 | 0.630 | 0.504 |
| num. syllables | -0.12 | 0.02 | -4.82 | <0.001 | <0.001 | -0.10 | 0.02 | -4.50 | <0.001 | <0.001 |
| stimulus order | 0.03 | 0.01 | 4.74 | <0.001 | <0.001 | 0.07 | 0.01 | 9.70 | <0.001 | <0.001 |
| motor rate * auditory rate | -0.04 | 0.02 | -2.60 | **0.013** | **0.009** | 0.01 | 0.02 | 0.37 | 0.765 | 0.709 |

Random effects

| | | High synchronizers | | Low synchronizers | |
|---|---|---|---|---|---|
| Group | Name | Variance | STD | Variance | STD |
| Trial-ID | (Intercept) | 0.213540 | 0.46210 | 0.189647 | 0.43548 |
| Subjects | Syllabic rate | 0.003219 | 0.01794 | 0.000547 | 0.02339 |
| Residual | | 0.300949 | 0.54859 | 0.288296 | 0.52693 |
| N (subs) | | 27 | | 20 | |
| Observations | | 8100 | | 6000 | |
| Marginal $R^2$ | | 0.471 | | 0.481 | |

**Supplementary Table 5. HG general model: Predicting speech tracking in HG.**

Predicted speech tracking in HG

| Predictors | beta | SE | t | $p_{FDR}$ | p |
|---|---|---|---|---|---|
| syllabic rate | -9.27 | 0.80 | -11.53 | **<0.001** | **<0.001** |
| syllabic rate$^2$ | 0.14 | 0.78 | 0.18 | 0.981 | 0.858 |
| HG | -0.04 | 0.07 | -0.65 | 0.720 | 0.514 |
| SMA-HG coupling | -0.06 | 0.06 | -1.03 | 0.626 | 0.305 |
| SMA | 0.22 | 0.07 | 3.01 | **0.023** | **0.003** |
| IFG-HG coupling | 0.00 | 0.06 | 0.02 | 0.984 | 0.984 |
| IFG | -0.11 | 0.08 | -1.50 | 0.429 | 0.134 |
| HG × SMA-HG coupling | -0.04 | 0.06 | -0.59 | 0.720 | 0.553 |
| HG × SMA | 0.11 | 0.07 | 1.56 | 0.429 | 0.119 |
| SMA x SMA-HG coupling | 0.07 | 0.08 | 0.90 | 0.626 | 0.366 |
| HG × IFG-HG coupling | 0.04 | 0.08 | 0.55 | 0.720 | 0.585 |
| HG × IFG | -0.07 | 0.09 | -0.86 | 0.626 | 0.391 |
| IFG x IFG-HG coupling | -0.08 | 0.07 | -1.14 | 0.626 | 0.254 |
| (HG × SMA) × SMA-HG coupling | 0.00 | 0.08 | -0.05 | 0.984 | 0.958 |
| (HG × IFG) × IFG-HG coupling | 0.19 | 0.08 | 2.39 | 0.094 | **0.018** |

Random effects

| Group | Name | Variance | STD |
|---|---|---|---|
| Subjects | Syllabic rate | 0.001271 | 0.03566 |
| Residual | | 0.659555 | 0.81213 |
| N (subs) | | 54 | |
| Observations | | 270 | |
| Marginal $R^2$ | | 0.396 | |

**Supplementary Table 6. Predicting speech tracking in HG, including behaviorally measured auditory-motor synchronization (PLV) as predictor.**

Predicted speech tracking in HG

| Predictors | beta | SE | t | $p_{FDR}$ | p |
|---|---|---|---|---|---|
| syllabic rate | -9.26 | 0.79 | -11.70 | **<0.001** | **<0.001** |
| syllabic rate$^2$ | 0.11 | 0.78 | 0.14 | 0.989 | 0.886 |
| HG | 0.01 | 0.07 | 0.07 | 0.989 | 0.942 |
| SMA-HG coupling | -0.04 | 0.06 | -0.75 | 0.668 | 0.457 |
| SMA | -0.18 | 0.15 | -1.20 | 0.398 | 0.230 |
| IFG-HG coupling | 0.03 | 0.06 | 0.40 | 0.817 | 0.688 |
| IFG | 0.19 | 0.16 | 1.21 | 0.298 | 0.229 |
| HG × SMA-HG coupling | -0.03 | 0.06 | -0.57 | 0.725 | **0.572** |
| HG × SMA | 0.12 | 0.07 | 1.71 | 0.240 | **0.076** |
| SMA x SMA-HG coupling | 0.05 | 0.08 | 0.65 | 0.698 | **0.514** |
| HG × IFG-HG coupling | 0.09 | 0.07 | 1.14 | 0.403 | **0.254** |
| HG × IFG | -0.11 | 0.09 | -1.27 | 0.398 | **0.206** |
| IFG x IFG-HG coupling | -0.09 | 0.07 | -1.39 | 0.391 | **0.164** |
| (HG × SMA) × SMA-HG coupling | 0.00 | 0.07 | -0.01 | 0.989 | 0.989 |
| (HG × IFG) × IFG-HG coupling | 0.19 | 0.08 | 2.44 | 0.098 | **0.016** |
| PLV | 0.58 | 0.34 | 1.71 | 0.241 | 0.076 |
| PLV x SMA | 0.77 | 0.29 | 2.62 | 0.089 | **0.009** |
| PLV x IFG | -0.62 | 0.33 | -1.90 | 0.224 | 0.059 |

Random effects

| Group | Name | Variance | STD |
|---|---|---|---|
| Subjects | Syllabic rate | 0.001026 | 0.03203 |
| Residual | | 0.653650 | 0.80849 |
| N (subs) | | 54 | |
| Observations | | 270 | |
| Marginal $R^2$ | | 0.415 | |

**Supplementary Table 7. Group models: Predicting speech tracking in HG, separately for high and low synchronizers.**

Predicted speech tracking in HG

| | High synchronizers | | | | | Low synchronizers | | | | |
|---|---|---|---|---|---|---|---|---|---|---|
| Predictors | beta | SE | t | $p_{FDR}$ | p | beta | SE | t | $p_{FDR}$ | p |
| syllabic rate | -6.56 | 0.81 | -8.15 | **<0.001** | **<0.001** | -6.03 | 0.75 | -8.04 | **<0.001** | **<0.001** |
| syllabic rate$^2$ | 0.14 | 0.72 | 0.20 | 0.892 | 0.843 | 0.36 | 0.74 | 0.48 | 0.779 | 0.663 |
| HG | 0.12 | 0.12 | 1.01 | 0.612 | 0.315 | -0.18 | 0.14 | -1.33 | 0.476 | 0.186 |
| SMA | 0.28 | 0.13 | 2.15 | 0.180 | **0.034** | 0.14 | 0.10 | 1.36 | 0.476 | 0.177 |
| SMA-HG coupling | -0.09 | 0.10 | -0.88 | 0.612 | 0.383 | -0.08 | 0.12 | -0.69 | 0.714 | 0.491 |
| IFG | -0.44 | 0.14 | -3.02 | **0.025** | **0.003** | -0.04 | 0.13 | -0.34 | 0.840 | 0.735 |
| IFG-HG coupling | 0.05 | 0.11 | 0.45 | 0.782 | 0.654 | 0.08 | 0.10 | 0.79 | 0.687 | 0.429 |
| HG × SMA | 0.29 | 0.14 | 2.01 | 0.187 | **0.047** | -0.09 | 0.10 | -0.92 | 0.640 | 0.360 |
| HG × SMA-HG coupling | 0.01 | 0.10 | 0.14 | 0.892 | 0.892 | 0.00 | 0.10 | 0.01 | 0.992 | 0.992 |
| SMA × SMA-HG coupling | -0.08 | 0.13 | -0.58 | 0.751 | 0.563 | 0.12 | 0.13 | 0.97 | 0.640 | 0.333 |
| HG × IFG | 0.12 | 0.14 | 0.88 | 0.612 | 0.381 | -0.69 | 0.18 | -3.90 | **0.002** | **<0.001** |
| HG × IFG-HG coupling | 0.08 | 0.12 | 0.72 | 0.691 | 0.475 | -0.09 | 0.16 | -0.57 | 0.760 | 0.570 |
| IFG × IFG-HG coupling | -0.17 | 0.13 | -1.28 | 0.543 | 0.204 | -0.02 | 0.09 | -0.20 | 0.900 | 0.844 |
| (HG × SMA) × SMA-HG coupling | -0.05 | 0.13 | -0.41 | 0.782 | 0.685 | 0.16 | 0.10 | 1.66 | 0.401 | 0.100 |
| (HG × IFG) × IFG-HG coupling | 0.11 | 0.10 | 1.14 | 0.582 | 0.254 | 0.22 | 0.18 | 1.27 | 0.476 | 0.208 |

Random effects

| | | High synchronizers | | Low synchronizers | |
|---|---|---|---|---|---|
| Group | Name | Variance | STD | Variance | STD |
| Subjects | Syllabic rate | 0.001891 | 0.04348 | 0.000 | 0.000 |
| Residual | | 0.492249 | 0.70160 | 0.8133 | 0.9018 |
| N (subs) | | 28 | | 20 | |
| Observations | | 140 | | 100 | |
| Marginal $R^2$ | | 0.557 | | 0.511 | |

**Supplementary Table 8. STG general model: Predicting speech tracking in STG.**

Predicted speech tracking in STG

| Predictors | beta | SE | t | $p_{FDR}$ | p |
|---|---|---|---|---|---|
| syllabic rate | -8.75 | 0.83 | -10.51 | **<0.001** | **<0.001** |
| syllabic rate$^2$ | 1.957 | 0.80 | 1.96 | 0.205 | 0.051 |
| STG | -0.18 | 0.06 | -3.24 | **0.011** | **0.001** |
| SMA-STG coupling | -0.05 | 0.05 | -0.86 | 0.690 | 0.388 |
| SMA | 0.11 | 0.06 | 1.85 | 0.208 | 0.065 |
| IFG-STG coupling | 0.12 | 0.06 | 2.08 | 0.205 | 0.039 |
| IFG | -0.01 | 0.06 | -0.18 | 0.947 | 0.859 |
| STG × SMA-STG coupling | -0.01 | 0.06 | -0.17 | 0.947 | 0.868 |
| STG × SMA | -0.07 | 0.06 | -1.11 | 0.607 | 0.269 |
| SMA × SMA-STG coupling | -0.06 | 0.06 | -1.03 | 0.607 | 0.304 |
| STG × IFG-STG coupling | -0.02 | 0.06 | -0.40 | 0.947 | 0.691 |
| STG × IFG | 0.00 | 0.07 | 0.07 | 0.947 | 0.947 |
| IFG × IFG-STG coupling | -0.07 | 0.06 | -1.16 | 0.607 | 0.247 |
| (STG × SMA) × SMA-STG coupling | 0.03 | 0.06 | 0.51 | 0.947 | 0.611 |
| (STG × IFG) × IFG-STG coupling | 0.01 | 0.07 | 0.18 | 0.947 | 0.857 |

Random effects

| Group | Name | Variance | STD |
|---|---|---|---|
| Subjects | Syllabic rate | 0.0005466 | 0.02338 |
| Residual | | 0.6000252 | 0.77461 |
| N (subs) | | 50 | |
| Observations | | 250 | |
| Marginal $R^2$ | | 0.388 | |

**Supplementary Table 9. Predicting speech tracking in STG, including behaviorally measured auditory-motor synchronization (PLV) as predictor.**

Predicted speech tracking in STG

| Predictors | beta | SE | t | $p_{FDR}$ | p |
|---|---|---|---|---|---|
| **syllabic rate** | -8.78 | 0.81 | -10.82 | **<0.001** | **<0.001** |
| syllabic rate$^2$ | 1.58 | 0.78 | 2.02 | 0.142 | **0.045** |
| **STG** | -0.19 | 0.06 | -3.41 | **0.005** | **0.011** |
| SMA-STG coupling | -0.03 | 0.05 | -0.52 | 0.722 | 0.604 |
| SMA | -0.11 | 0.15 | -0.77 | 0.702 | 0.444 |
| IFG-STG coupling | 0.44 | 0.14 | -0.52 | 0.722 | **0.032** |
| **IFG** | 0.44 | 0.14 | 3.23 | **0.007** | **0.001** |
| STG × SMA-STG coupling | -0.02 | 0.06 | -0.40 | 0.730 | 0.692 |
| STG × SMA | -0.05 | 0.07 | -0.78 | 0.702 | 0.436 |
| SMA x SMA-STG coupling | -0.08 | 0.06 | -1.41 | 0.338 | 0.160 |
| STG × IFG-STG coupling | -0.03 | 0.06 | -1.63 | 0.281 | 0.646 |
| STG × IFG | -0.00 | 0.07 | -0.00 | 1.000 | 1.000 |
| IFG x IFG-STG coupling | -0.10 | 0.06 | -1.63 | 0.281 | 0.104 |
| (STG × SMA) × SMA-STG coupling | 0.06 | 0.06 | 0.84 | 0.702 | 0.383 |
| (STG × IFG) × IFG-STG coupling | 0.03 | 0.07 | 0.87 | 0.722 | 0.633 |
| PLV | 0.14 | 0.28 | -0.51 | 0.722 | 0.610 |
| PLV x SMA | 0.43 | 0.27 | 1.57 | 0.281 | 0.118 |
| **PLV x IFG** | -1.01 | 0.28 | -3.64 | **0.003** | **<0.001** |

Random effects

| Group | Name | Variance | STD |
|---|---|---|---|
| Subjects | Syllabic rate | 0.000818 | 0.0286 |
| Residual | | 0.589251 | 0.7676 |
| N (subs) | | 50 | |
| Observations | | 250 | |
| Marginal $R^2$ | | 0.421 | |

**Supplementary Table 10. Group models: Predicting speech tracking in STG, separately for high and low synchronizers.**

Predicted speech tracking in STG

| | High synchronizers | | | | Low synchronizers | | | | |
|---|---|---|---|---|---|---|---|---|---|
| Predictors | beta | SE | t | $p_{FDR}$ | beta | SE | t | $p_{FDR}$ | p |
| syllabic rate | -6.50 | 0.77 | -8.43 | **<0.001** | -5.57 | 0.81 | -6.88 | **<0.001** | **<0.001** |
| syllabic rate$^2$ | 1.53 | 0.74 | 2.09 | 0.128 | 0.87 | 0.73 | 1.19 | 0.508 | 0.237 |
| STG | -0.16 | 0.09 | -1.91 | 0.135 | -0.49 | 0.14 | -3.43 | **0.008** | **0.001** |
| SMA | 0.26 | 0.09 | 3.05 | **0.015** | 0.21 | 0.13 | 1.58 | 0.477 | 0.119 |
| SMA-STG coupling | 0.06 | 0.08 | 0.81 | 0.519 | -0.14 | 0.11 | -1.34 | 0.504 | 0.183 |
| IFG | -0.15 | 0.08 | -1.91 | 0.135 | 0.30 | 0.13 | 2.24 | 0.149 | **0.028** |
| IFG-STG coupling | 0.10 | 0.08 | 1.30 | 0.390 | 0.10 | 0.15 | 0.70 | 0.555 | 0.486 |
| STG × SMA | -0.23 | 0.09 | -2.57 | **0.046** | 0.23 | 0.19 | 1.19 | 0.504 | 0.238 |
| STG × SMA-STG coupling | -0.05 | 0.09 | -0.48 | 0.670 | 0.09 | 0.12 | 0.71 | 0.555 | 0.477 |
| SMA × SMA-STG coupling | -0.07 | 0.09 | -0.74 | 0.527 | -0.12 | 0.12 | -1.00 | 0.521 | 0.321 |
| STG × IFG | -0.10 | 0.10 | -0.95 | 0.519 | -0.13 | 0.18 | -0.71 | 0.555 | 0.482 |
| STG × IFG-STG coupling | -0.03 | 0.08 | -0.43 | 0.670 | 0.16 | 0.16 | 0.99 | 0.521 | 0.326 |
| IFG × IFG-STG coupling | -0.24 | 0.08 | -3.05 | **0.015** | -0.04 | 0.13 | -0.29 | 0.778 | 0.771 |
| (STG × SMA) × SMA-STG coupling | 0.08 | 0.10 | 0.82 | 0.519 | 0.04 | 0.13 | 0.28 | 0.778 | 0.778 |
| (STG × IFG) × IFG-STG coupling | 0.10 | 0.10 | 1.04 | 0.519 | 0.13 | 0.14 | 0.90 | 0.536 | 0.368 |

Random effects

| | | High synchronizers | | Low synchronizers |
|---|---|---|---|---|
| Group | Name | Variance | STD | |
| Subjects | (Intercept) | 0.03455 | 0.1859 | |
| Residual | | 0.51855 | 0.7201 | |
| N (subs) | | 27 | | |
| Observations | | 135 | | |
| Marginal $R^2$ | | 0.484 | | |

| Group | Name | | Variance | STD |
|---|---|---|---|---|
| Subjects | Syllabic rate | | 0.000 | 0.000 |
| Residual | | | 0.6032 | 0.7767 |
| N (subs) | | | 19 | |
| Observations | | | 95 | |
| Marginal $R^2$ | | | 0.542 | |
